# DNA Break-Induced Epigenetic Drift as a Cause of Mammalian Aging

**DOI:** 10.1101/808659

**Authors:** Motoshi Hayano, Jae-Hyun Yang, Michael S. Bonkowski, Joao A. Amorim, Jaime M. Ross, Giuseppe Coppotelli, Patrick T. Griffin, Yap Ching Chew, Wei Guo, Xiaojing Yang, Daniel L. Vera, Elias L. Salfati, Abhirup Das, Sachin Thakur, Alice E. Kane, Sarah J. Mitchell, Yasuaki Mohri, Emi K. Nishimura, Laura Schaevitz, Neha Garg, Ana-Maria Balta, Meghan A. Rego, Meredith Gregory-Ksander, Tatjana C. Jakobs, Lei Zhong, Hiroko Wakimoto, Raul Mostoslavsky, Amy J. Wagers, Kazuo Tsubota, Stephen J. Bonasera, Carlos M. Palmeira, Jonathan G. Seidman, Christine E. Seidman, Norman S. Wolf, Jill A. Kreiling, John M. Sedivy, George F. Murphy, Philipp Oberdoerffer, Bruce R. Ksander, Luis A. Rajman, David A. Sinclair

**Affiliations:** Paul F. Glenn Center for Biology of Aging Research, Department of Genetics, Blavatnik Institute, Harvard Medical School, Boston, MA 02115, USA; Department of Ophthalmology, Keio University School of Medicine, 35 Shinanomachi, Shinjuku-ku, Tokyo, 160-8582, Japan; IIIUC-Institute of Interdisciplinary Research, University of Coimbra, Coimbra, Portugal; Zymo Research Corporation, Irvine, CA 92614, USA; Laboratory for Ageing Research, Department of Pharmacology, School of Medical Sciences, The University of New South Wales, Sydney, NSW 2052, Australia; Experimental Gerontology Section, Translational Gerontology Branch, National Institute on Aging, NIH, Baltimore, MD 21224, USA; Department of Stem Cell Biology, Medical Research Institute, Tokyo Medical and Dental University, 1-5-45 Yushima, Bunkyo-ku, Tokyo, 113-8510, Japan; Vium Inc., San Mateo, CA 94402 USA; Schepens Eye Research Institute, Department of Ophthalmology, Massachusetts Eye and Ear Infirmary, Harvard Medical School, Boston, MA 02114; The Massachusetts General Hospital Cancer Center, Harvard Medical School, Boston, MA 02114, USA; Department of Genetics, Harvard Medical School, Boston, MA 02115, USA; Paul F. Glenn Center for the Biology of Aging Research and Department of Stem Cell and Regenerative Biology, Harvard Stem Cell Institute, Harvard University, Cambridge, MA, USA; Division of Geriatrics, Department of Internal Medicine, University of Nebraska Medical Center, 3028 Durham Research Center II, Omaha, NE, 68198-5039, USA; Department of Life Sciences, Faculty of Sciences and Technology, University of Coimbra, Coimbra, Portugal; Department of Pathology, University of Washington, Seattle, WA 98195, USA; Department of Molecular Biology, Cell Biology and Biochemistry, Brown University, Providence, RI 02912, USA; Program in Dermatopathology, Department of Pathology, Brigham & Women’s Hospital, Harvard Medical School, Boston, MA 02115, USA; Laboratory of Receptor Biology and Gene Expression, National Cancer Institute, NIH, Bethesda, MD 20892, USA; Department of Integrative Structural and Computational Biology, The Scripps Research Institute, La Jolla, CA, USA

## Abstract

There are numerous hallmarks of aging in mammals, but no unifying cause has been identified. In budding yeast, aging is associated with a loss of epigenetic information that occurs in response to genome instability, particularly DNA double-strand breaks (DSBs). Mammals also undergo predictable epigenetic changes with age, including alterations to DNA methylation patterns that serve as epigenetic “age” clocks, but what drives these changes is not known. Using a transgenic mouse system called “ICE” (for inducible <u>c</u>hanges to the <u>e</u>pigenome), we show that a tissue’s response to non-mutagenic DSBs reorganizes the epigenome and accelerates physiological, cognitive, and molecular changes normally seen in older mice, including advancement of the epigenetic clock. These findings implicate DSB-induced epigenetic drift as a conserved cause of aging from yeast to mammals.

**One Sentence Summary:** DNA breaks induce epigenomic changes that accelerate the aging clock in mammals

## INTRODUCTION

For cells to maintain optimal function they must counteract the entropic loss of both genetic and epigenetic information. They achieve this, at least for some length of time, because they are open systems, utilizing energy from their environment to maintain genetic and epigenetic information, including the DNA sequence, transcriptional networks, chromatin modifications, and DNA methylation (DNAme) patterns (Kane and Sinclair, 2019; Keller, 2009).

In the 1950s, Szilard and Medawar independently proposed that aging is caused by a loss of genetic information (Medawar, 1952; Szilard, 1959). Since then, numerous studies have evidence in support of this model. For example, cells from old individuals exposed to the environment accumulate abundant mutations (Martincorena et al., 2018; Yizhak et al., 2019) and organisms with defects in DNA repair appear to undergo aspects of accelerated aging, as exemplified by segmental progeroid disorders such as trichothiodystrophy, and Werner, Rothmund-Thompson and Cockayne syndromes, (Carrero et al., 2016; Harkema et al., 2016; Park et al., 1999; Sinclair et al., 1997).

The type of DNA damage that is most consistently linked to aging is the DSB, a particularly toxic form of DNA damage that occurs at a rate of 10 to 50 per cell per day (Vilenchik and Knudson, 2003). A recent comparison of 18 different rodent species, from mice to beavers, found that, of all DNA repair processes, the repair of DSBs was by far most highly correlated with lifespan (Tian et al., 2019). Indeed, in mice, defects in DSB repair due to the deletion of *Ku70*, *Ku80* or *Ercc1* result in progeroid phenotypes (Li et al., 2007; Niedernhofer et al., 2006) while overexpression of the murine SIRT6 gene, a DSB repair protein that declines with age, extends the lifespan of mice (Kaya et al., 2015; Mao et al., 2011).

In recent years, however, studies have questioned the importance of mutations in aging. The number of mutations in old yeast and in the tissues of old mice are relatively rare, though there are exceptions, such as mammalian cells exposed to the environment or when clones amplify in the hematopoietic system (Dolle et al., 1997; Kaya et al., 2015). In addition, some engineered strains of mice with high levels of free radicals or mutation rates show little to no sign of premature aging or exhibit a shortened lifespan (Narayanan et al., 1997) while mice with defects in DNA repair, such as *Ercc2* (*Xpd*)^m/m^, exhibit a progeroid phenotype, ostensibly without any increase in the mutation rate (Dolle et al., 2006). Perhaps the strongest argument against a loss of genomic information as a major cause of aging is the fact that it is possible to clone animals, such as dogs, goats, sheep, almost all of which have normal, healthy lifespans (Burgstaller and Brem, 2017).

In the 1990s, studies of *Saccharomyces cerevisiae* indicated that yeast aging is caused by a loss of epigenetic, rather than genetic, information (Kennedy et al., 1997; Sinclair et al., 1997). In old yeast cells, relocalization of the NAD^+^- and CR-responsive silent information regulator complex (Sir2/3/4) from silent mating type loci to the nucleolus is believed to result in the simultaneous expression of both **a** and *α* mating type information, resulting in sterility, a hallmark of old yeast cells (Smeal et al., 1996). During yeast aging, global epigenetic changes are also seen, including decreases in heterochromatin, epigenetic regulators, altered histone modifications (e.g. H3K56ac and H4K16ac), histone occupancy, and an increase in gene transcription globally. The fact that the modulation of chromatin factors extends yeast lifespan – such as overexpression of *SIR2* or histone genes, or deletion of the histone acetyltransferase gene *HAT2* or histone methyltransferase gene *SET2* – also indicates that epigenetic changes are a cause of yeast aging, not merely a biomarker (Dang et al., 2009; Feser et al., 2010; Hu et al., 2014; Kaeberlein et al., 1999; Rosaleny et al., 2005; Ryu et al., 2014).

Multicellular organisms also experience changes to chromatin and the epigenome during aging, examples of which include global decreases in DNA methylation and a loss of the imbalance between euchromatin (H3K4me3) and heterochromatin marks (H3K9me3 and H3K27me3) (Benayoun et al., 2015; Pal and Tyler, 2016; Sen et al., 2016). In worms, for example, deletion of genes for the H3K4 trimethylation complex that catalyzes the formation of euchromatin extends lifespan (Greer et al., 2010; Greer et al., 2011). In flies, levels of repressive heterochromatin marks, such as H3K9me2, H3K9me3 and H3K27me3 are altered during aging (Larson et al., 2012; Ma et al., 2018; Wood et al., 2010), the suppression of which by calorie restriction or overexpression of fly dSir2 maintains a youthful epigenome, suppresses the age-dependent expression of potentially deleterious transposable elements, and extends lifespan (Jiang et al., 2013; Rogina and Helfand, 2004; Wood et al., 2016). The long-lived naked mole rat has a relatively stable epigenome, with increased levels of repressive marks compared to mice (Tan et al., 2017) and in mice and humans, a loss of histones is seen in senescent cells (Ivanov et al., 2013; O’Sullivan et al., 2010) and in quiescent muscle stem cells (satellite cells) during aging (Liu et al., 2013).

Some of these changes are remarkably consistent between individuals and species. For example, the methylation state of select CpG sites can be used to accurately predict the biological age and eventual lifespan of individuals of a wide range of mammalian species, and of individuals of different species, arguing the underlying mechanism is conserved (Hannum et al., 2013; Horvath, 2013; Petkovich et al., 2017; Weidner et al., 2014).

Together, these findings have led to a shift from viewing aging as a random process to one that is non-random and potentially driven by reproducible and predictable epigenetic changes. But this shift begs new questions. What causes the mammalian epigenome to change over time and how does it impact aging? And if DNA damage has a role in aging, how can these seemingly random events produce similar changes between individuals, making it appear as though it were a program?

Again, clues have come from *S. cerevisiae*. A major driver of epigenetic change in yeast is the DSB, which triggers a damage signal that recruits epigenetic regulators away from gene promoters to the DNA break site where they assist with repair, relieving repression at repetitive loci such as rDNA and telomeres. Examples of such dual-role factors include Sir2, Hst1, Rpd3, Gcn5, and Esa1 (Martin et al., 1999; McAinsh et al., 1999; Mills et al., 1999; Tamburini and Tyler, 2005). The “Relocalization of Chromatin Modifiers” or “RCM” hypothesis, a component of the “Information Theory of Aging” (Kane and Sinclair, 2019; Lu et al., 2019), proposes that this phenomenon evolved as an ancient response to DNA damage that turns on stress-response genes to enhance survival and, in the case of yeast, prevents mating until the damage is repaired because it would likely be lethal (Oberdoerffer and Sinclair, 2007). According to the model, most DNA repair factors return to their original sites on the genome after DSB repair is complete, turning off the stress response, but not completely. With repeated DNA damage responses, the epigenomic landscape is progressively altered to the point where cells remain in a chronically stressed state, which eventually disrupts their cellular identity.

There is now abundant evidence that RCM occurs in higher organisms (Kane and Sinclair, 2019). Like Sir2 in yeast, three of the seven mammalian Sir2 homologs, SIRT1, SIRT6 and SIRT7, have dual roles as epigenetic regulators and facilitators of DNA break repair (Mao et al., 2011; Oberdoerffer et al., 2008; Paredes et al., 2018). Together, these histone deacetylases form repressive chromatin at telomeres, transposons, rDNA, and hundreds of genes involved in metabolism, circadian rhythms, inflammation, and DNA repair, among others (Kawahara et al., 2011; Oberdoerffer et al., 2008; Paredes et al., 2018; Simon et al., 2019; Van Meter et al., 2014). In response to an ATM-mediated DNA damage signal, RCM proteins relocalize to DSB sites to assist with repair by modifying histones and recruiting other DNA repair proteins (Dobbin et al., 2013; Mao et al., 2011; Oberdoerffer et al., 2008; Toiber et al., 2013). After repair, the original chromatin and epigenetic status of the cells is reestablished although not completely, thereby introducing informational noise (O’Hagan et al., 2008). Over time there is a cumulative shift away from the original epigenetic state (Kim et al., 2018). Consistent with the RCM hypothesis, overexpression of SIRT1 or SIRT6 increases genome stability, suppresses retrotransposons, delays transcriptional changes with age (Oberdoerffer et al., 2008; Roichman et al., 2016) and, in the case of SIRT6, extends lifespan (Kanfi et al., 2012).

Together, these data point to a cause of aging in eukaryotes being the dysregulation of transcriptional networks and epigenetic information over time, driven by a conserved mechanism that evolved to co-regulate genetic and epigenetic responses to adversity (Mills et al., 1999; Oberdoerffer et al., 2008). This idea is consistent with antagonistic pleiotropy, whereby a biological system that benefits the survival of young individuals can be deleterious later in life when the forces of natural selection no longer prevail (Williams, 1957).

To test cause and effect, specifically whether DNA breaks induce epigenetic changes that accelerate aging in a mammal, we generated a transgenic mouse called “ICE” (for Inducible <u>C</u>hanges to the <u>E</u>pigenome). We find that disruption of the epigenome in a young mouse induces aging at the molecular, histological, and physiological level, including an acceleration of the DNA methylation “epigenetic clock.” These data, combined with that of our accompanying paper indicating that cellular aging is associated with a smoothening of the chromatin landscape and a loss of cell identity (see Yang et al., co-submitted manuscript), are consistent with the hypothesis that the loss of epigenetic information is an upstream cause of mammalian aging.

## RESULTS

### The ICE System Creates Transient and Non-Mutagenic DSBs

To test the hypothesis that DSBs induce epigenetic changes that contribute to aging, we designed and created a murine system that allows for precise temporal and spatial control over DSBs. The DSBs are created primarily in non-coding regions at frequencies only a few-fold above normal background levels. To create non-mutagenic DSBs we employed I-*Ppo*I, a homing endonuclease encoded by a selfish genetic element in the slime mold *Physarum polycephalum*, which has been used previously to study DSB repair in the context of the RCM response, including the roles of ATM, NBS1, SIRT1, SIRT6, HDAC1, and LSD1 (Berkovich et al., 2007; Dobbin et al., 2013; McCord et al., 2009; Mosammaparast et al., 2013).

I-*Ppo*I recognizes the DNA sequence CTCTCTTAA▼GGTAGC (Monnat et al., 1999), which occurs 20 times in the mouse genome, 19 of which are non-coding, and none of which occur in mitochondrial DNA (Berkovich et al., 2007) (**Figure S1A and S1B**). The actual number of sites that are cut is far lower than that, presumably due to limited DNA accessibility. Unlike other methods of creating DSBs, such as CRISPR, chemicals and radiation, I-*Ppo*I creates “sticky DNA ends” that are repaired without inducing a strong DNA damage response or a mutation (Yang et al., co-submitted manuscript).

The transgenic system consists of two components (**Figure 1A**): the first is a fusion of the I-*Ppo*I gene to the C-terminus of a tamoxifen (TAM)-regulated mutant estrogen receptor domain gene (ER^T2^) and a transcriptional *loxP*-STOP-*loxP* cassette that is targeted by Cre recombinase (Berkovich et al., 2007; Kim et al., 2016). The second is a tamoxifen-regulated Cre recombinase gene (Cre-ER^T2^) under the control of the human ubiquitin C promoter to ensure whole-body expression (Ruzankina et al., 2007). When the two components are combined *in vivo* and tamoxifen is provided, Cre-ER^T2^ is expressed and directed to the nucleus where it excises the stop cassette, facilitating transcription of the ER^T2^-HA-I-*Ppo*I-IRES-GFP gene cassette. The protein product, ER^T2^-I-*Ppo*I, is then directed to the nucleus. Upon removal of tamoxifen, ER^T2^-I-*Ppo*I is no longer directed to the nucleus and is degraded in the cytoplasm. C57BL6/J transgenic mice containing each of these components were crossed, generating Het ER^T2^-I-*Ppo*I and Cre-ER^T2^, named Inducible <u>C</u>hanges to the <u>E</u>pigenome or “ICE” mice, with wildtype (WT), I-*Ppo*I and Cre mice serving as negative controls (**Figure 1B**).

**Figure 1.**
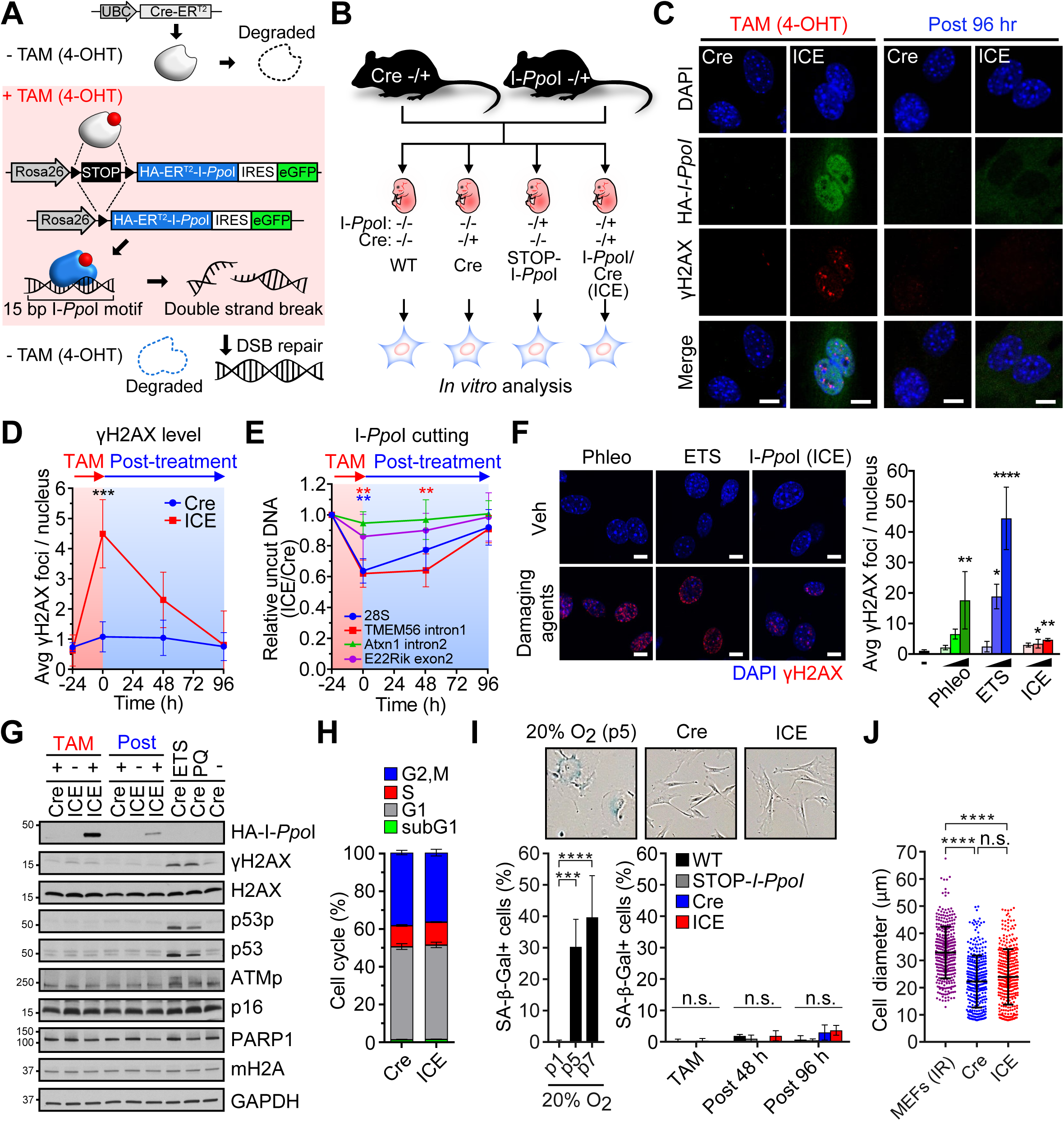
A Cell-Based System to Create Inducible Changes to the Epigenome (ICE) (A) Schematic of the ICE system based on a tamoxifen-inducible I-*Ppo*I endonuclease. (B) Breeding scheme to generate ICE mice and negative controls (WT, Cre and I-*Ppo*I). (C and D) γH2AX foci in DAPI-stained nuclei of MEFs from ICE mice and Cre controls after tamoxifen (4-OHT, 0.5 µM) induction. Scale bar, 10 µm. Two-way ANOVA-Bonferroni. (E) qPCR analysis of DNA cutting at I-*Ppo*I canonical sites. One-way ANOVA-Bonferroni. (F) DNA damage response induced by I-*Ppo*I (4-OHT, 0.1, 0.5, 1 µM) vs. other DNA damaging agents, etoposide (ETS, 1, 10, 25 µM) and phleomycin (Phleo, 1, 25, 50 µg/ml). Scale bar, 10 µM. One-way ANOVA-Bonferroni. (G) Western blot of proteins that are involved in and downstream of the DNA damage response. Blots assessing p53p and γH2AX were reprobed for p53 and H2AX using antibodies raised in different species. (H) Cell cycle profile in Cre and ICE cells 96 hrs post-tamoxifen treatment. (I) Percentage of senescence-associated β-galactosidase positive (blue) cells during and after tamoxifen treatment compared to replicative senescent cells. p, passage. One-way ANOVA-Bonferroni. (J) Cell diameter after recovery from I-*Ppo*I induction vs. irradiated (senescent) cells. One-way ANOVA-Bonferroni. Data are mean (n≥3) ± SD. n.s.: p > 0.05; *p < 0.05; **p < 0.01; ***p< 0.001; **** p < 0.0001.

To generate an equivalent, cell-based ICE system, mouse embryonic fibroblasts (MEFs) were isolated from Iittermates at Day E13.5 and cultured under low oxygen conditions (3% v/v). After the addition of tamoxifen (0.5 μM 4-OHT), HA-I-*Ppo*I was detected in the nucleus (**Figure 1C, 1D, S1C and S1D**) and the number of serine 139-phosphorylated H2AX (γH2AX) foci, a marker of DSBs, reached a maximum of 4-fold above background at the 24 hour time point, with the extent of cutting locus-dependent (**Figure 1E, S1E and S1F**). Compared to the DNA damaging agent etoposide (a topoisomerase II inhibitor) and phleomycin (a free-radical inducer), the number of γH2AX foci, the extent of DNA breakage, and the DNA damage response in the ICE cells was minimal (**Figure 1F-J and S1G**). Over a 24 hr time course of I-*Ppo*I induction, there were no detectable changes in cell cycle profile, senescence, or number of aneuploid cells (**Figure 1H-1J and S1H**). At the 28S rDNA, there was no change in mutation frequency, RNA levels, or overall translation efficiency in the treated ICE cells (**Figure S2A-S2H)**.

### The ICE System Induces Non-Mutagenic Cuts *In Vivo*

To test the ICE system *in vivo*, we performed whole-body I-*Ppo*I induction for three weeks in 4-6 month-old mice by providing a modified AIN-93G purified rodent diet containing tamoxifen citrate (360 mg/kg) and monitored the mice for another 10 months. Unless stated specified, all experiments followed this experimental timeline (**Figure 2A**). The extent of STOP cassette removal was similar in muscle (67%), liver (71%), hippocampus (61%) and cortex (72%) (**Figure 2B**) and HA-I-*Ppo*I, γH2AX and eGFP were detectable during tamoxifen treatment in all tissues tested (**Figure 2C and 2D**).

**Figure 2.**
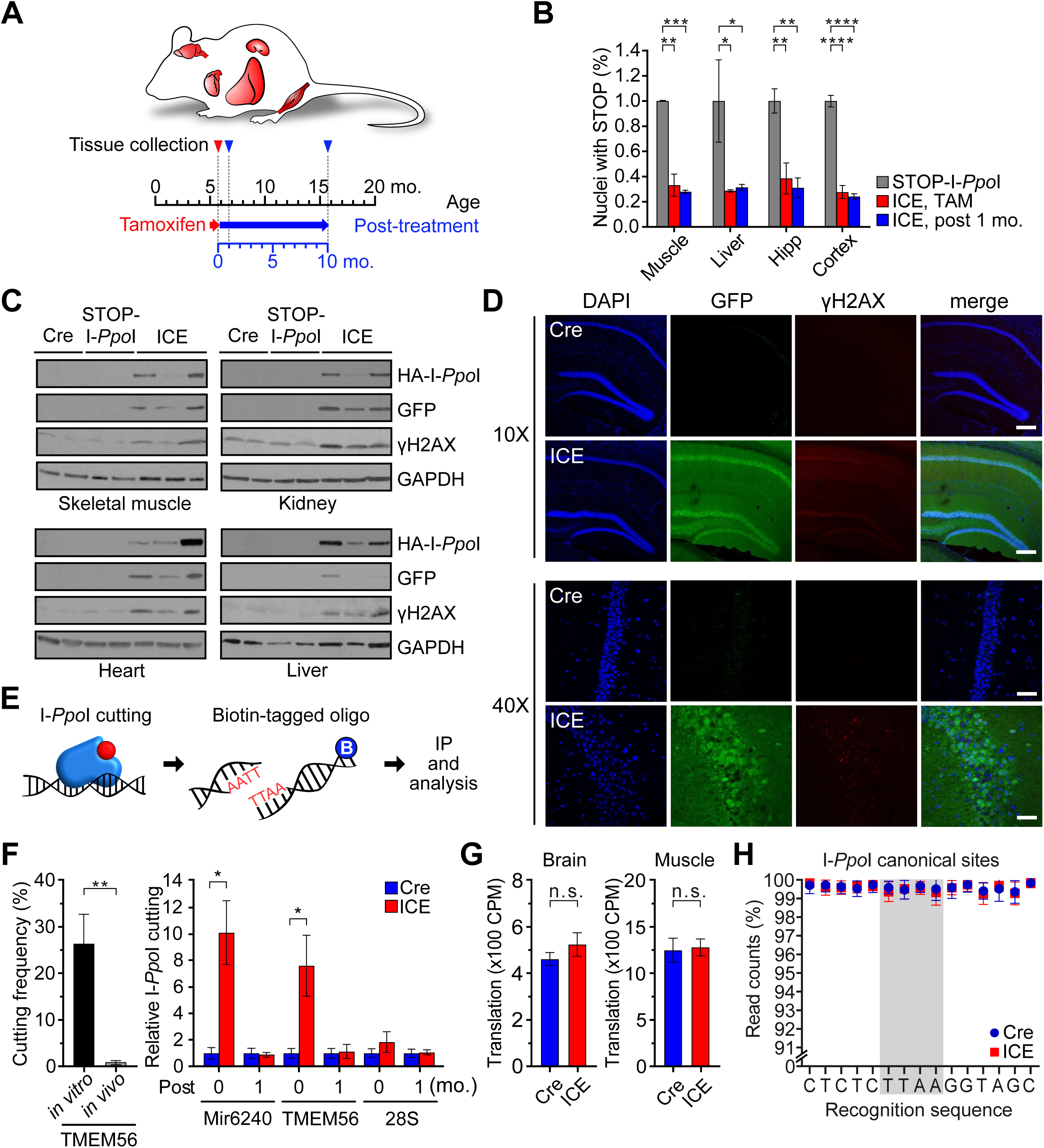
Validation of the ICE Mouse. (A) Timeline of the induction of I-*Ppo*I and phenotypic assessment of mice. (B) Removal of the transcriptional STOP cassette in major tissues. One-way ANOVA-Bonferroni. (C) Western blot of tissues probed with anti-HA to detect I-*Ppo*I expression and γH2AX after a 4 week tamoxifen treatment. (D) Hippocampal sections immunostained for GFP (as a proxy for IRES-linked I-*Ppo*I expression) and γH2AX. Scale bar, 200 µm (10X) and 50 µm (40X). (E and F) Immunoprecipitation and quantification of I-*Ppo*I cut sites in skeletal muscle during and after tamoxifen treatment (0-vs 1-month post-treatment). Two-tailed Student’s *t* test. (G) Protein translation in brain and skeletal muscle assessed by metabolic ^35^S-labelling. Two-tailed Student’s *t* test. (H) Whole genome sequencing (>50x) of the 19 canonical I-*Ppo*I recognition sites in skeletal muscle 1 month after ceasing I-*Ppo*I induction. Data are mean (n≥3) ± SEM. n.s.: p > 0.05; *p < 0.05; **p < 0.01; ***p< 0.001; **** p < 0.0001.

To assess the location and extent of I-*Ppo*I cutting, we adapted an end-capture qPCR assay (Chailleux et al., 2014) that uses a biotinylated oligo harboring the overhang 5‘-TTAA-3’ to precipitate DNA ends resulting from a I-*Ppo*I cut (**Figure 2E**). In skeletal muscle, a tissue well studied during aging, the Tmem56 intron site (Chr3:121249934-121249948) was cut ∼25-fold less efficiently compared to purified DNA and was undetectable 1-month after tamoxifen was removed. Interestingly, cutting at the 28S site was undetectable during and after treatment, presumably due to inaccessibility (**Figure 2F**), and there was no change in 28S rRNA levels or overall protein synthesis (**Figure 2G, S3A and S3B**). Telomere length was not significantly different between Cre and ICE tissues 10-month post-treatment (**Figure S3C**) and no differences in mutation frequency were detected in gastrocnemius muscle by whole genome sequencing 1-month post-treatment, either at canonical or non-canonical I-*Ppo*I recognition sites or across the genome (Wittmayer et al., 1998) (**Figure 2H, S3D and S3E**). Together, these data show that the ICE system is able to induce specific DNA breaks that are completely repaired and cause no overt effects on the mice during tamoxifen induction (Yang et al., co-submitted manuscript).

### Non-Mutagenic DNA Cutting Phenocopies Age-Associated Physiological Changes

As mice age, they undergo a characteristic physical and physiological changes, including alopecia, hair greying, kyphosis, decreased body weight, decreased motion in the dark phase, and reduced respiration during the day as they utilize fat rather than carbohydrate as an energy source (Ackert-Bicknell et al., 2015; Harkema et al., 2016; Houtkooper et al., 2011; Koks et al., 2016). If the RCM hypothesis has validity, a short period of induction of I-*Ppo*I should introduce epigenomic noise and accelerate some, if not all, of these changes. To test this, we placed cohorts of 4-6 month-old ICE mice on a tamoxifen-containing diet for three weeks and assessed them for another 10 months (**Figure 3A**), with identically treated WT, Cre, and I-*Ppo*I littermate controls serving as negative controls. During the three-weeks of I-*Ppo*I induction, there were no detectable differences between ICE mice and controls in terms of behavior, activity, or food intake. After one month, however, there were visual differences in the ICE mice compared to the controls, such as slight alopecia and loss of pigment on the feet, tail, ears and nose common features of middle-aged WT mice (Liu et al., 2019; Nishimura et al., 2005) (**Figure 3B and S4K**).

**Figure 3.**
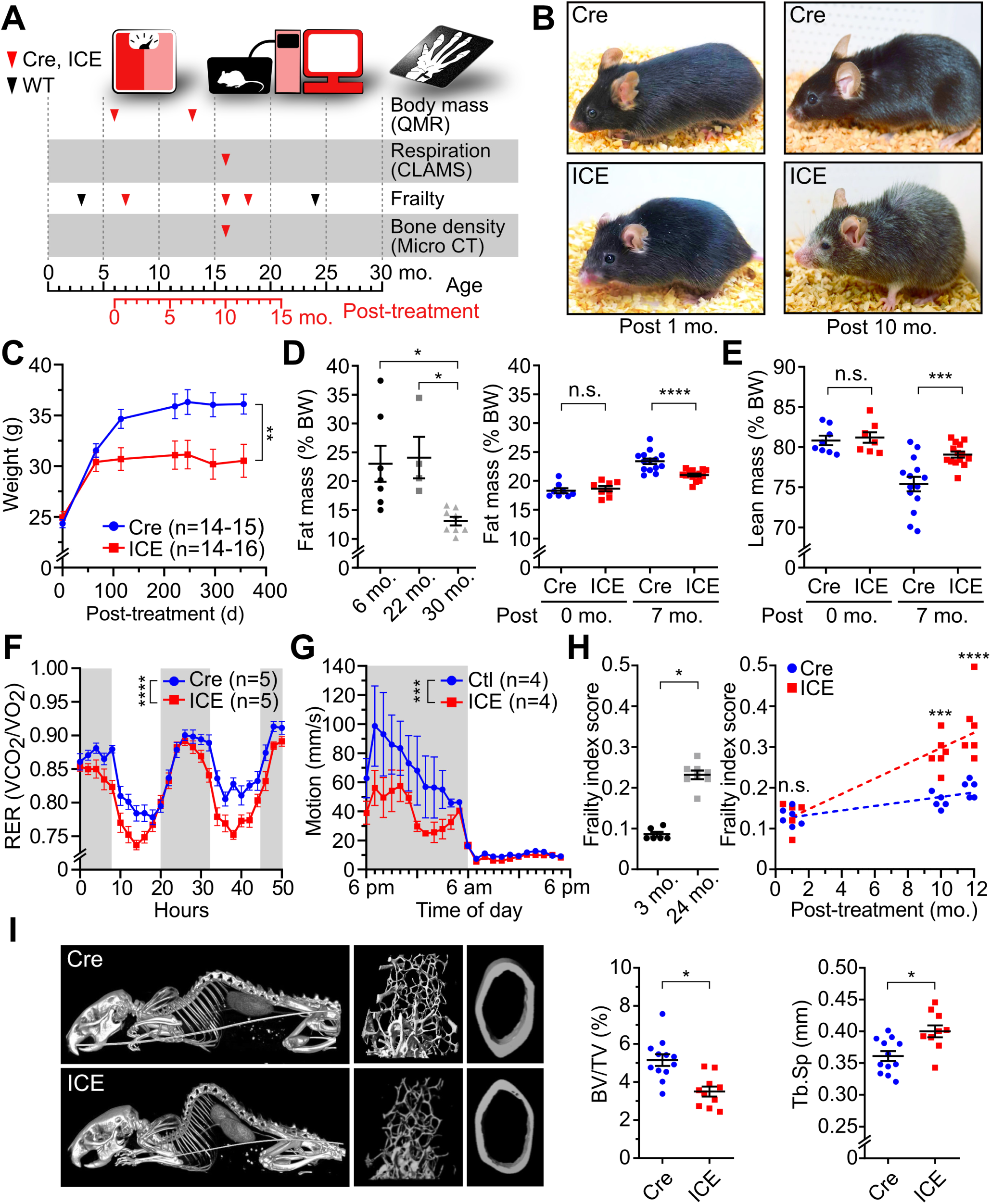
ICE Mice Are an Accelerated Phenocopy of Normal Aging. (A) Timeline of phenotypic assessments of mice. (B) Representative images of Cre and ICE mice. (C-E) Weight and body mass of Cre and ICE mice. Repeated measures one-way ANOVA (C). One-way ANOVA-Bonferroni (D, left). Two-way ANOVA-Bonferroni (D right and E). (F) Respiratory Exchange Rate (RER) of Cre and ICE mice. Repeated measures one-way ANOVA. (G) Average activity of Cre and ICE mice for 55 days. Repeated measures one-way ANOVA. (H) Frailty index scores of Cre, ICE, wild type 3 and 24 month-old mice. Two-tailed Student’s *t* test (left) or two-way ANOVA-Bonferroni (right). (I) CT of whole skeleton and micro-CT of trabecular and cortical bones of Cre and ICE mice. Kyphosis assessment (left), bone/tissue volume (center) and trabecular separation (right). Two-tailed Student’s *t* test. Data are mean ± SEM. n.s.: p > 0.05; *p < 0.05; **p < 0.01; ***p< 0.001; **** p < 0.0001.

By the 10-month post-treatment time-point, all the ICE mice, but none of the controls, exhibited classic features of old age, including reduced body weight and fat mass independent of food intake (**Figure 3B, 3C-3E and S4A-S4H**), a lower respiratory exchange ratio (RER) during the day (**Figure 3F and S4I**) and decreased motion in the dark phase (**Figure 3G**).

To provide a longitudinal, quantitative measure of health, we utilized the frailty index (FI), which combines 31 parameters, such as body weight, temperature, coat condition, grip strength, mobility, vision and hearing (Whitehead et al., 2013). To establish a baseline, we established the FI scores of WT mice between 3 and 24 months of age, noting an increase of 2.5-fold. There was also no significant difference in FI scores between ICE mice and controls at the 1-month post-treatment timepoint. By the 10- and 12-month timepoints, however, the FI scores of the ICE mice were substantially higher than controls, closer to 24 month-old WT mice (p=0.0006 and <0.0001, respectively) (**Figure 3H**), along with accelerated kyphosis and a loss of cortical bone thickness and trabecular bone density in the inner layer (Ferguson et al., 2003; Katzman et al., 2010), both common features of aging (**Figure 3I**).

### ICE Mice Exhibit Age-Related Physiological Changes to Skin and Eye

One of the hallmarks of age-related physical decline in mice and in humans is a loss of visual acuity causes by an increase in lens opacity and the degeneration of retinal function including the loss of retinal ganglion cells (RGCs) and their axons (Calkins, 2013; Samuel et al., 2011; Wolf et al., 2000). Ten months after tamoxifen treatment, there was significantly more lens opacity in the ICE mice compared to Cre controls (**Figure 4A, 4B and S4J**). RGC cell bodies, which reside in the innermost retinal layer and their axons form the nerve fiber layer that congregates in the posterior of the eye (**Figure 4C**), are particularly vulnerable to age-related stress when they pass through the optic nerve head (Downs, 2015). ICE mice had significantly fewer optic nerve axons in the myelinated region, paralleling what is typically seen in 24 month-old wild type mice (**Figure 4D**).

**Figure 4.**
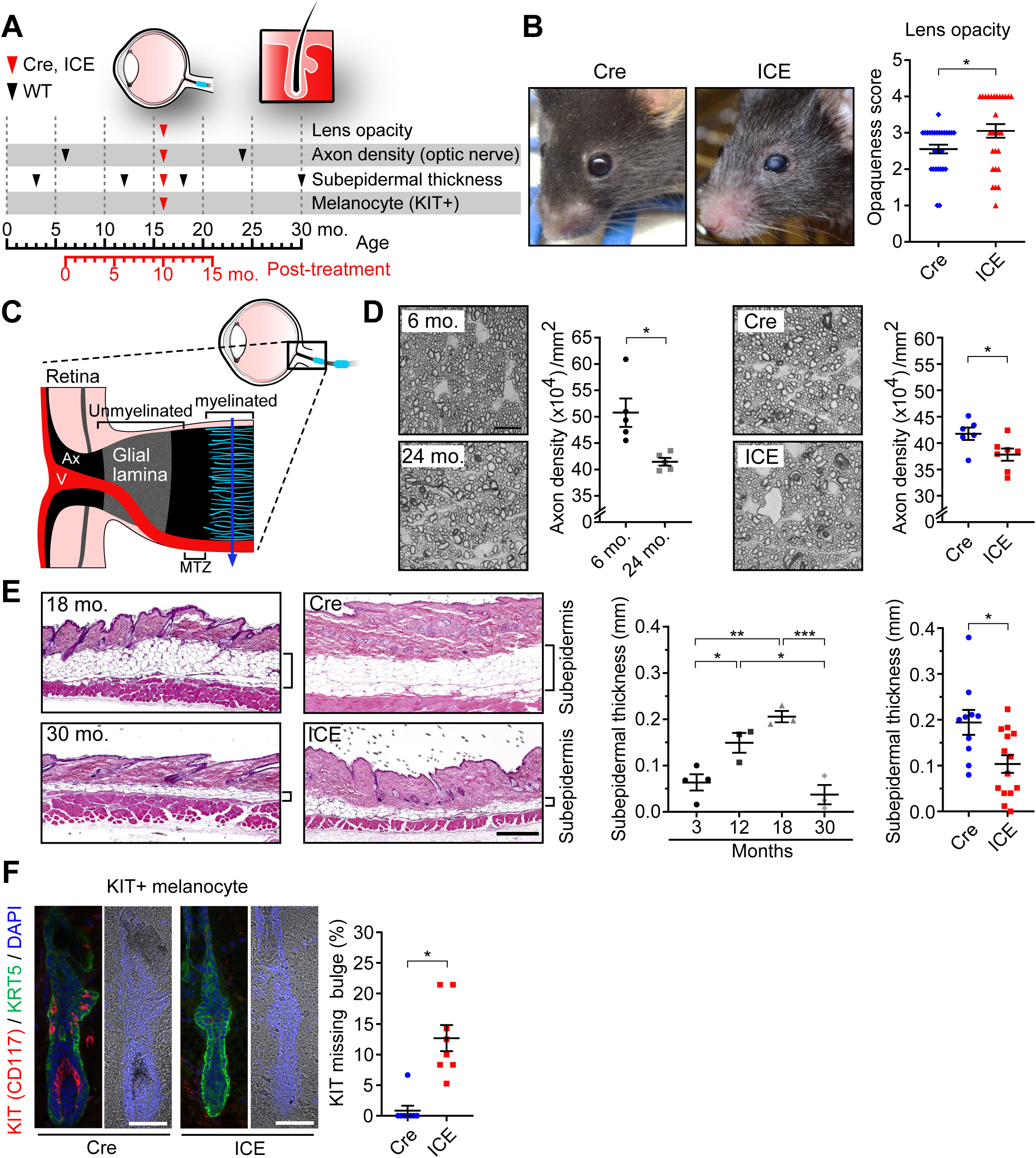
Progeroid Features of the Skin and Eyes of ICE Mice. (A) Timeline of phenotype assessments. (B) Representative images and quantification of lens opacity. Two-tailed Student’s *t* test. (C) Schematic diagram of the optic nerve head indicating the location of tissues obtained for axon counts (solid line). V, retinal blood vessels; MTZ, myelination transition zone; Ax, axon bundles. (D) Representative photomicrographs of PPD stained myelinated optic nerve axons. Scale bar, 10 μm. Quantification of healthy axons, represented as axon density (x10^4^)/mm^2^. Two-tailed Student’s *t* test. (E) H&E staining of subcutaneous fat layers and subepidermal thickness of back skin from old WT, Cre and ICE mice. Scale bar, 500 µm. One-way ANOVA-Bonferroni (middle) or two-tailed Student’s *t* test (right). (F) KIT (CD117), KRT55 and DAPI staining of back skin and percent of hair follicle bulges without c-kit staining. Scale bar, 50 μm. Two-tailed Student’s *t* test. (G) 5-hydroxymethyl-cytosine (5hmC) staining of back skin of WT, Cre and ICE mice. Scale bar, 10 µm. One-way ANOVA-Bonferroni (middle) or two-tailed Student’s *t* test (right). Data are mean ± SEM. n.s.: p > 0.05; *p < 0.05; **p < 0.01; ***p< 0.001; **** p < 0.0001.

Mammalian skin also progresses through well-defined changes during aging. Subepidermal thickness typically increases until middle age then declines rapidly, along with hair greying due to a loss of KIT/CD117-positive melanocyte stem cells (Gomes et al., 2013; Matsumura et al., 2016; Nishimura et al., 2005). These features were accelerated in the ICE mice, including hair greying and a loss of both subepidermal thickness and KIT/CD117 positive melanocytes (**Figure 4E and 4F**).

### ICE Mice Exhibit Age-Related Physiological Changes to Muscle

Age-related changes to skeletal muscle have been well characterized and include decreases in exercise endurance, muscle strength, mass, vascularization and mitochondrial function (Das et al., 2019; Demontis et al., 2013) (**Figure 5A**). Ten months after treatment, ICE mice had significantly less muscle mass compared to controls (**Figure 5B**) and exhibited a decrease in running endurance and buildup of lactate equivalent to WT mice at 30 months of age (**Figure 5C and 5D**). The limb grip strength of ICE mice was also lower than age-matched Cre controls (**Figure 5E**), more similar to mice at an older age. Key hallmarks of muscle aging at the cellular level include decreases in ATP levels, mitochondrial DNA copy number, increased mitochondrial area and alterations in subsarcolemmal and intermyofibrillar mitochondrial morphology (Demontis et al., 2013; Leduc-Gaudet et al., 2015). The ICE mice but not controls displayed all of these changes at the 10-month post-treatment time point, resembling 20 to 24 month-old wild type mice (**Figure 5F-5J and S5A-S5D**).

**Figure 5.**
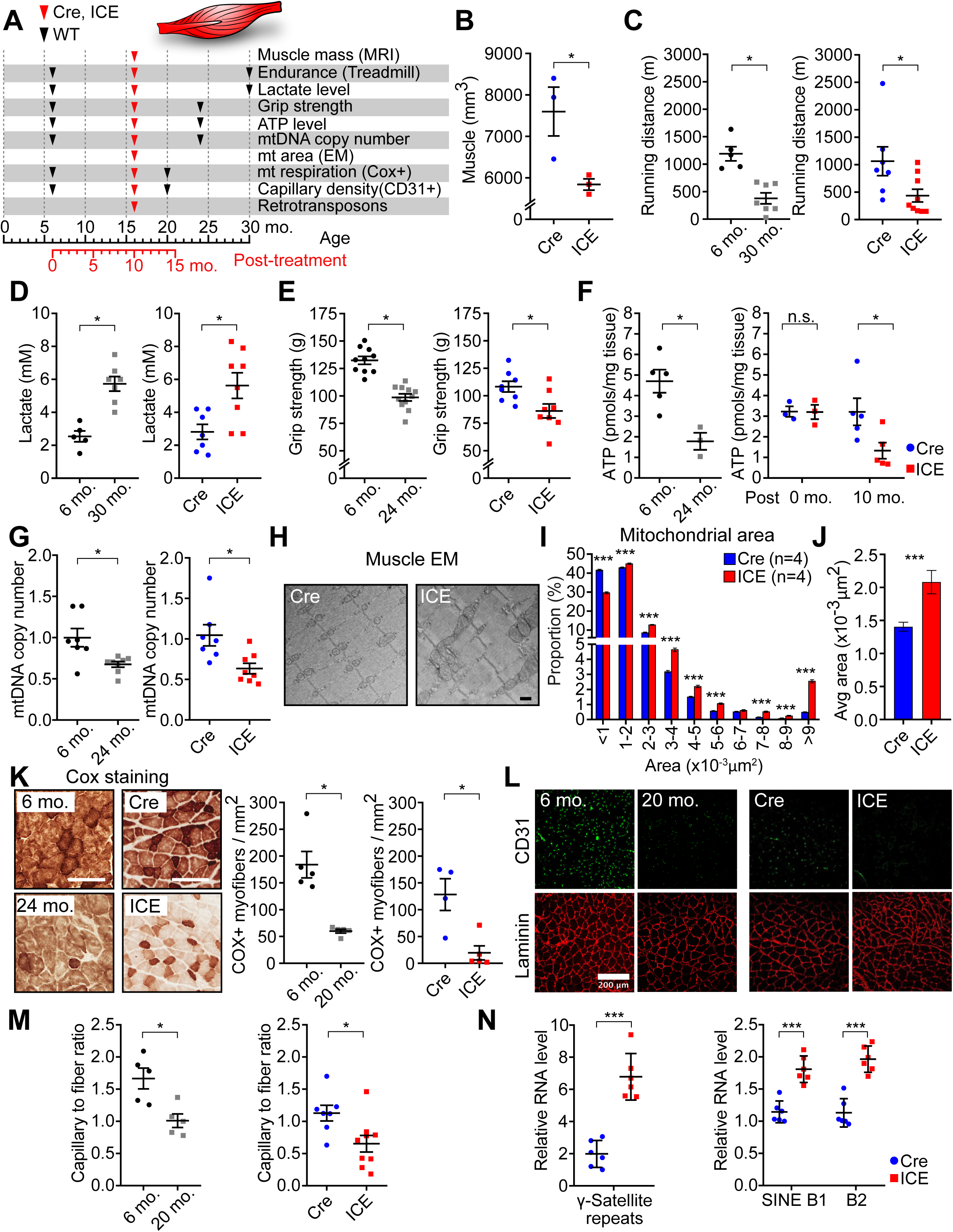
Progeroid Features of ICE Mouse Muscles. (A) Timeline of phenotypic assessments of mice. (B) Muscle mass assessed by MRI. Two-tailed Student’s *t* test. (C-D) Treadmill endurance and blood lactate build-up after exercise in WT, Cre and ICE mice. Two-tailed Student’s *t* test. (E) Grip strength measured as maximal “peak force“. Two-tailed Student’s *t* test. (F) ATP levels in gastrocnemius muscle. Two-tailed Student’s *t* test (let) or two-way ANOVA-Bonferroni (right). (G-J) Mitochondrial DNA copy number, morphology and area. Scale bar, 500 nm. Two-tailed Student’s *t* test. (K) Cytochrome oxidase (COX) staining of gastrocnemius muscle. Scale bar, 100 µm. Two-tailed Student’s *t* test. (L and M) Gastrocnemius immunostained with laminin (red) and CD31(green), markers of the extracellular matrix and capillaries, respectively, and the ratio thereof. Two-tailed Student’s *t* test. (N) Quantification of RNA from repetitive DNA elements in gastrocnemius muscle. Two-tailed Student’s *t* test. Data are mean ± SEM. n.s.: p > 0.05; *p < 0.05; **p < 0.01; ***p< 0.001.

Another well-known hallmark of muscle aging is a decrease in the abundance of cytochrome oxidase, a component of complex IV in the OXPHOS system (Wenz et al., 2009). The ICE mice at 16 months of age had 6-fold fewer COX-positive myofibers, paralleling the difference between 6 and 24 month-old WT mice (**Figure 5K**). Consistent with this, the abundance of H3K27ac, a marker of active promoters and enhancers, was lost from genes involved in metabolism (Yang et al., co-submitted manuscript).

One of the most obvious changes during aging in mammals is loss of muscle microvasculature (Das et al., 2019). The capillary to fiber ratio in skeletal muscles of 16 month-old ICE mice was about half that of the Cre mice and more similar to ratios seen in 24 month-old mice (**Figure 5L and 5M**). Loss of silencing at repetitive elements and retrotransposons is also a conserved feature of aging and a potential driver of inflammation (De Cecco et al., 2019; Oberdoerffer et al., 2008) and this was also seen in the skeletal muscle of ICE mice compared to Cre controls (**Figure 5N, S5E and S5F**). On average, ICE mice had a thinner left ventricular (LV) posterior wall than Cre controls, implying possible dilated cardiomyopathy, with no difference in LV internal diameter or ejection fraction (**Figure S5G**).

### ICE Mice Experience Changes to the Brain That Are Typically Those Seen in Much Older Mice

One of the main hallmarks of mammalian aging is a decline in the function of the CNS, manifested as defects in motor coordination and cognition (**Figure 6A**) (Johnson et al., 2018; Ungvari et al., 2017). In the dark phases, ambulatory activity was 50% lower in the ICE mice relative to Cre controls (**Figure 6B**) and gait coordination, based on swing and stance times, was also impaired (**Figure S6A-S6C**). The hippocampus is critical for spatial and memory consolidation, the function of which declines predictably with age (Gallagher et al., 2010; Miller and O’Callaghan, 2005; Park and Reuter-Lorenz, 2009). In the fear-conditioning paradigm, short-term memory is measured by placing them in specific contexts and subjecting them to an electric shock on Day 1, which they should recall on Day 2 and exhibit a freezing response. Because these parameters have not been well characterized for aged C57BL/6J, we established a baseline. The immediate freezing response was similar between young and old mice (6- vs. 24- and 30-month) on Day 1, but on the second day, 75% of the young mice compared to <40% of old mice froze, indicating the old mice had a reduced ability to recall the context from the day before (**Figure 6C-E)**. A very similar difference was seen when comparing Cre and ICE mice at 16 months of age, with >40% of the Cre controls responding to the context on the second day, compared to only about 24% of the ICE mice (**Figure 6D-E**).

**Figure 6.**
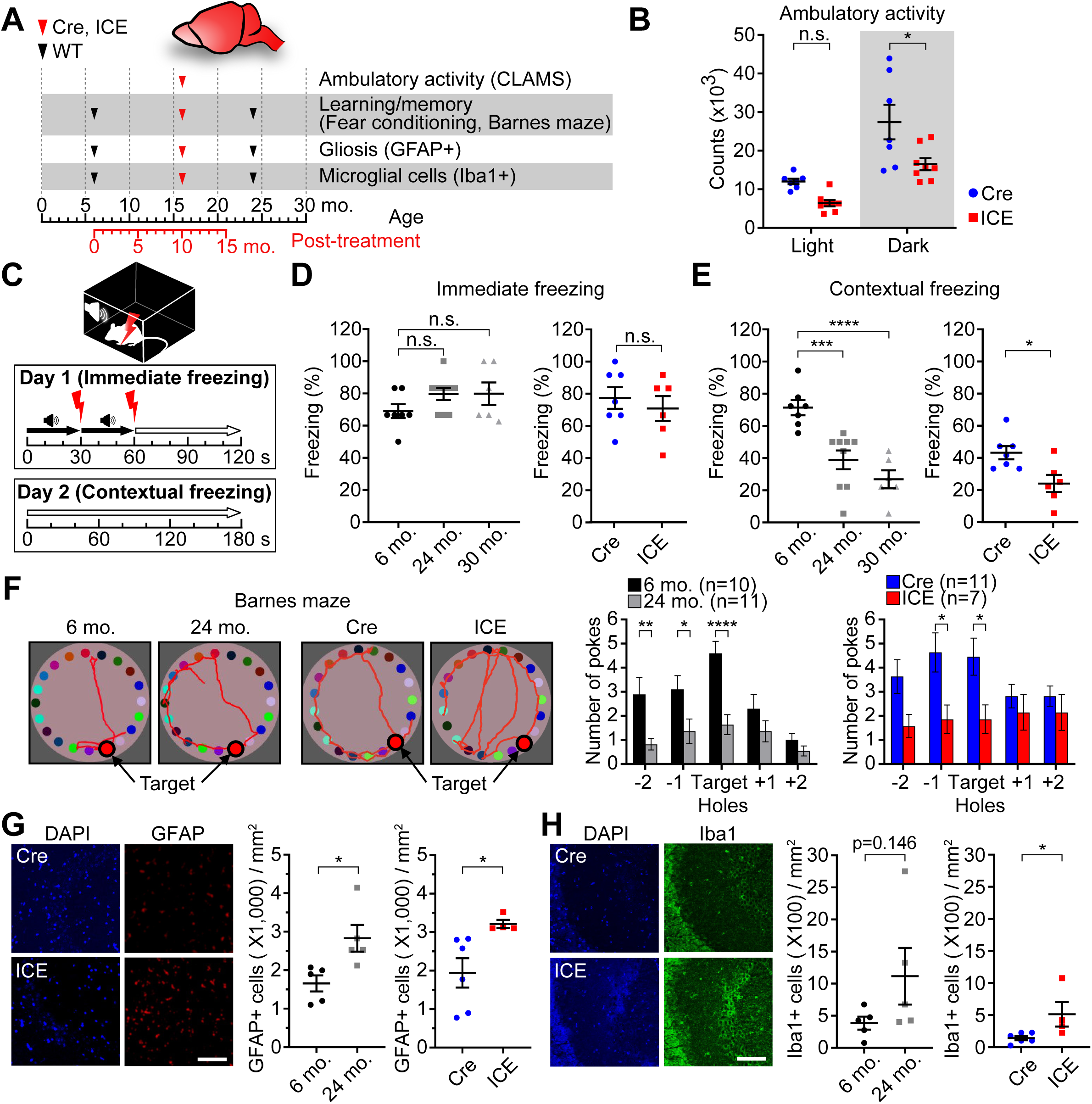
The Brains of ICE Mice Appear to Undergo Accelerated Aging. (A) Timeline of phenotypic assessments of mice. (B) Ambulatory activity in light and dark cycles. Two-way ANOVA-Bonferroni. (C and E) Immediate and contextual freezing in fear conditioning tests. One-way ANOVA-Bonferroni (D, left and E, left) or two-tailed Student’s *t* test (D, right and E, right). (F) Representative images of Barnes maze tests and mean number of pokes at each hole. Two-way ANOVA-Bonferroni. (G and H) Immunofluorescence of the hippocampal CA3 region immunostained for inflammatory markers GFAP and Iba1. Scale bar, 100 µm. Two-tailed Student’s *t* test. Data are mean ± SEM. n.s.: p > 0.05; *p < 0.05; **p < 0.01; ***p< 0.001; **** p < 0.0001.

Another measure of hippocampal function and long-term memory is the Barnes maze test. Over five days, mice learn to identify the location of a hiding box then, 7 days later, mice are re-tested for their ability to recall the location of the hiding box. The recall of ICE mice was about half that of age matched Cre controls, similar to the recall of 24 month-old WT mice (**Figure 6F**).

Within the central nervous system, astrocytes and microglia are critical mediators of the innate immune response. As mammals age, the innate immune system becomes hyper-activated and the number of activated microglia and astrocytes, which can be detected by staining for Iba1 and GFAP, increases (Baruch et al., 2014; Norden and Godbout, 2013). In the WT cohort, the hippocampi of 24 month-old mice had more GFAP- and Iba1-positive cells compared to 6-month-old mice, consistent with previous reports (Baruch et al., 2014). Paralleling normal aging, the ICE mice at 16 months of age had significant numbers of GFAP- and Iba1-positive cells, 1.6- and 3.5-fold greater than the Cre controls (**Figure 6G and 6H**). Together, these data indicate that ICE mice experience an acceleration of brain inflammation and memory loss, reminiscent of normal aging.

### ICE Mice Experience Age-Related Transcriptional Changes and Acceleration of the Epigenetic Clock

To provide a more quantitative assessment of biological age, we compared gene expression patterns and DNA methylation (DNAme) patterns of ICE and controls. In skeletal muscle, genes that were significantly dysregulated in ICE mice were positively correlated with changes in wild type 24 month-old mice (**Figure 7B-D, S7A, S7B and Table S2**). Notable examples include the mediator of p53-mediated cellular senescence *Cdkn1a* (Cyclin Dependent Kinase Inhibitor 1A or p21) (Beggs et al., 2004; Choudhury et al., 2007; Welle et al., 2004), *Myl4* (Myosin light chain 4), which encodes an embryonic form of myosin that is upregulated in aged mouse muscle (Lin et al., 2018), *Nlrc5* (NLR family CARD domain containing 5), which inhibits NF-κB activation and is one of the most significantly hypomethylated genes in centenarians (Zeng et al., 2018), and *Mrpl55* (mitochondrial ribosomal protein L55), which encodes a 39S mitochondrial ribosomal gene whose methylation status is associated with life expectancy (Weidner et al., 2014; Zhang et al., 2017).

**Figure 7.**
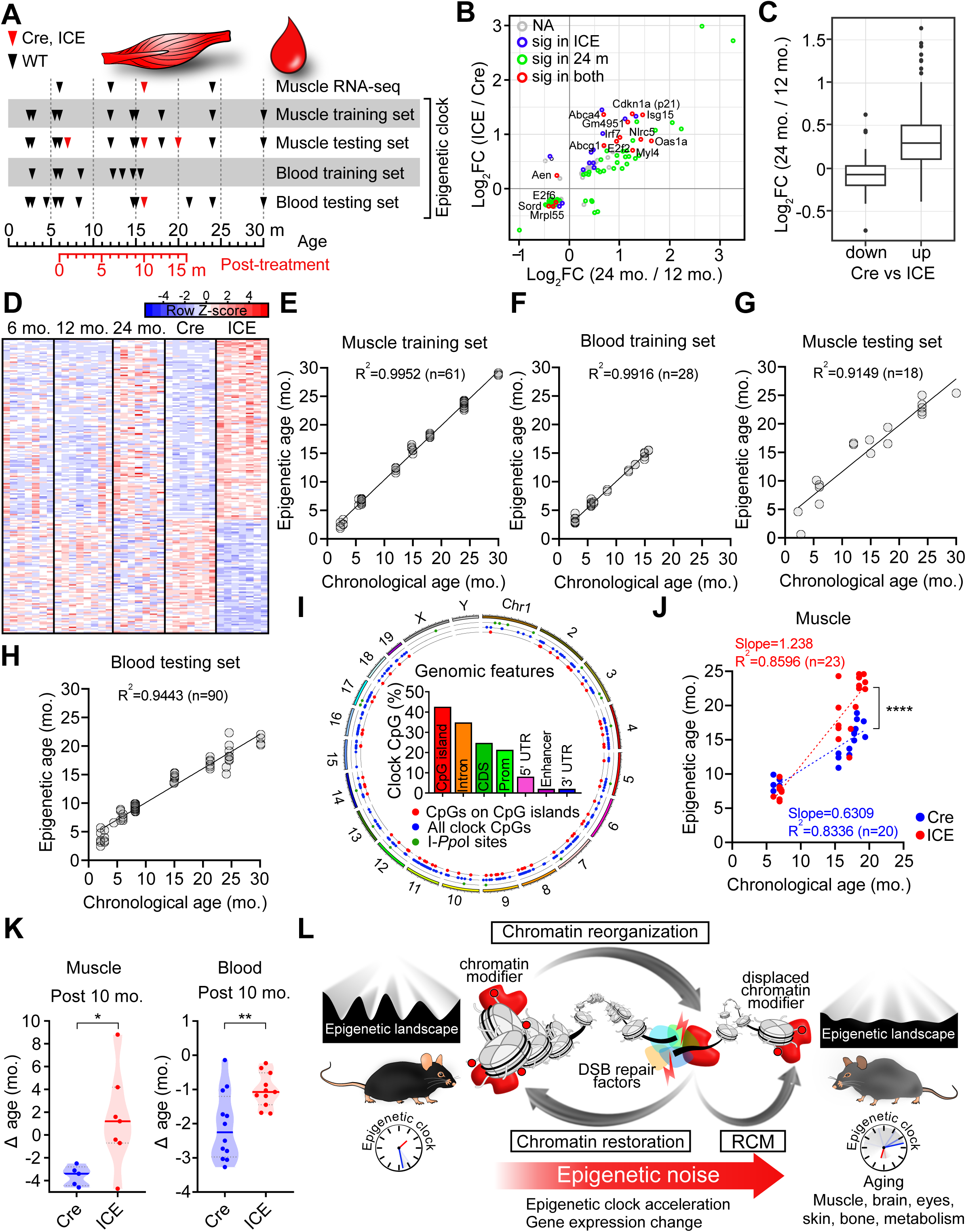
ICE Mice Experience an Acceleration of The Epigenetic Clock. (A) Timeline of phenotypic assessments of mice. (B) Scatter plot of genes significantly changed (p<0.01) in 10-month post-treated ICE mice and wild type 24 month-old mice. (C) Fold change of altered genes (padj<0.05, Cre vs ICE mice) in wild type 24 month-old mice. (D) Heatmaps of the top 200 most significantly altered genes in skeletal muscle of Cre and ICE mice. (E-H) Muscle and blood training (E and F) and testing sets (G and H) of the clock CpG sites in WT C57BL/6J mice. (I) Circos plot of genomic locations of I-*Ppo*I cut sites (green), clock GpG sites in CpG islands (red), and all clock sites (blue). (J) Epigenetic age of gastrocnemii of Cre and ICE mice 1-, 10-, and 14-month post-treatment. Linear regression analysis and Spearman correlation. (K) Epigenetic age of muscle and blood of Cre and ICE mice at 10-month post-treatment (Δ age = epigenetic age – chronological age.). Two-tailed Student’s *t* test. (J) Model for how DNA breaks and the resulting epigenetic noise contribute to aging. Data are mean ± SEM. *p < 0.05; **p < 0.01; **** p < 0.0001.

DNA methylation (DNAme) clocks are a robust indicator of biological age in mammals (Hannum et al., 2013; Horvath, 2013; Petkovich et al., 2017; Weidner et al., 2014). To assess the relative DNAme age of the ICE mice, we developed DNAme clocks for skeletal muscle and for blood by assessing WT mice at 8-10 different ages (**Figure 7A**). Reduced-representation bisulfite sequencing (RRBS) on 79 skeletal muscle and 118 whole blood samples identified 915 age-associated CpG loci for a blood panel and 2,048 CpG loci for a multi-tissue panel. Targeted bisulfite-sequencing libraries using a simplified whole-panel amplification reaction method (SWARM) was then used to assess DNA methylation changes at >2,000 CpG sites sequenced to >2,500x coverage. We used 61 WT muscle and 29 WT blood samples from male and female C57BL/6 mice aged 2 to 30 months to define the training set (**Figure 7E-F**) and selected 2,048 multi-tissue clock CpGs by ElasticNet regression model with CpG sites with at least 300 reads in all samples. The epigenetic age for muscle and blood samples was calculated as: epigenetic age *= inverse.F(b_0_ + b_1_CpG_1_+ ⋯ +b_n_CpG_n_)*, where the *b* are coefficients obtained from the trained model with *b_0_* being the intercept and *CpG* are methylation values of the loci.

Within the training dataset, the epigenetic age derived from the weighted sum of DNA methylation levels of age-modulated CpG sites was highly correlated with the chronological age of the individual samples, with R^2^ = 0.995 and 0.992 for the muscle and the blood clock, respectively (**Figure 7E-F and Table S3**). For validation, 18 muscle and 90 blood samples aged 2 to 30 months were used for the testing dataset (**Figure 7G-H**). The clocks performed well in the validation data sets with R^2^ = 0.915 and 0.944 for the muscle and blood clocks respectively, indicating that both were able to accurately estimate age using two completely independent datasets. Across the genome, 40% of the DNAme clock sites were in CpG islands, 30% were in introns, 20% were in coding sequences and less than 5% were in enhancers or 3’ UTRs. There was no overlap between the DNAme sites and known I-*Ppo*I recognition sequences (**Figure 7I**).

Using these two mouse clocks, the rate of epigenetic aging was estimated to be about 50% faster in ICE mice than in Cre controls (p<0.0001) (**Figure 7J)**. By subtracting chronological age from epigenetic age, we determined the change in age (Δ age). The difference between chronological and epigenetic age (delta age) also indicated the ICE mice were older than Cre controls for both the muscle and the blood clock (**Figure 7K**).

## DISCUSSION

During embryogenesis, eukaryotic cells establish their identity by laying down a specific pattern of epigenetic information (Waddington, 1957). Why and how mammals lose epigenetic information over time, and whether this process is an underlying cause of aging, are currently matters of speculation. Together, the results in this study, combined with those in the accompanying manuscript (Yang et al.), indicate that the induction of a mild DNA damage response accelerates epigenetic drift that causes phenotypes and molecular changes that resemble normal aging (**Figure 7L**). The data strongly argue that the process of DSB repair, even if it doesn’t lead to a mutation, alters the epigenome and accelerates aging in mice at the physiological, histological, and molecular levels, including an acceleration of the epigenetic clock.

As far as we are aware, these studies identify the first molecular driver of epigenetic change during aging and the first set for convincing evidence that they drive the aging process. Our accompanying paper (Yang et al.) adds further weight to our conclusion by demonstrating that non-mutagenic DNA breaks alter the structure of the epigenome in predictable ways that closely resemble normal aging, including histone modifications, DNA compartmentalization, a smoothening of the epigenetic landscape, and a loss of cellular identity. Previous findings in yeast and these new findings in mammals, lend support to the hypothesis that epigenetic drift driven by DSB repair is a conserved cause of aging in all eukaryotes.

Traditionally, the process of DNA damage checkpoint activation and DNA repair has been studied using mutagens or radiation doses that cause DNA damage substantially above background levels. The ICE system allows us to create DSBs at levels at more natural levels, only a few-fold above background, thus avoiding overt DNA damage signaling response, cell cycle arrest, aneuploidy, mutations, or cellular senescence (Yang et al., co-submitted manuscript). Initially, no profound changes at the physiological or molecular levels were observed in the treated ICE mice. Indeed, the epigenetic clock did not initially advance. Over the course of the next 10 months, however, every tissue we examined had deteriorated and developed signs of aging. This observation suggests that molecular changes occurring during or shortly after the treatment trigger an advancement of the epigenetic clock many months later. We don’t yet know what these triggers are, but we hypothesize that might be alterations in DNA, chromatin, or transcriptional networks that initiate a feed-forward cascade of deleterious events. Future work will be aimed at identifying these cascades.

In humans, there is an abundance of evidence linking DNA damage to aging, including cancer chemotherapy, radiation exposure, smoking, and progeroid diseases such as Werner and Cockayne’s syndrome (Hofstatter et al., 2018; Horvath and Levine, 2015; Maccormick, 2006; Nance and Berry, 1992; Salk et al., 1985). Similarly, in model organisms, deficiencies in DNA repair, such as Ercc1, BubR1, Ku70, and Xpd mutant mice, also appear to accelerate aspects of aging (Carrero et al., 2016; White and Vijg, 2016). But mutation accumulation as a main cause of aging has been hard to reconcile with other findings that nuclear mutations in old individuals are not only rarer than would be expected, they can occur with high frequency without causing signs of aging (Dolle et al., 2006; Dolle et al., 1997; Narayanan et al., 1997).

Studies in simple organisms such as yeast and flies have indicated that changes to the epigenome are a cause of aging (Imai and Kitano, 1998; Jiang et al., 2013; Mills et al., 1999; Oberdoerffer et al., 2008). By causing DNA damage without causing mutations in cells and in mice, we provide strong evidence that it is the cell’s reaction to DNA breaks, rather than actual mutations, that drives the aging clock. This idea is particularly appealing because it explains why aging proceeds through a predictable series of molecular and physiological changes, despite the fact that DNA damage can occur anywhere in the genome.

These data also help explain why DSB repair efficiency correlates with longevity in diverse species, but not other types of DNA repair such as NER and BER (Brown and Stuart, 2007; Tian et al., 2019). In our model, DSBs are a special type of damage that potently induces epigenetic change. Because transcription-coupled DNA repair (TCR) defects in ERCC1 mutant mice also mimic aging (Niedernhofer et al., 2006), it will be interesting to determine whether TCR also induces an RCM response, epigenomic changes, and acceleration of the DNAme clock. Given that ERCC1 plays an indispensable role in the repair of DSBs containing DNA secondary structures, including AT-rich DNA sequences at fragile sites and G-quadruplexes (Li et al., 2019), perhaps the fundamental underlying mechanism of the ICE and ERCC1 mutant phenotype is the same.

Individuals treated with DSB-generating agents, such as chemotherapy, X-rays, and gamma radiation, are known to experience an acceleration of aging (Garrett-Bakelman et al., 2019; Maccormick, 2006; Richardson, 2009). In this study, low levels of DSBs were surprisingly impactful, even months later, raising the specter that very low doses radiation and even DNA editing enzymes may have deleterious long-term effects on the epigenome and tissue function.

The negative impact of DSBs on the epigenome raises the question why RCM evolved in the first place. We hypothesize it is an ancient system that places cells in a state of high-alert while DSB repair is carried out. At the molecular levels, DSBs induce relocalization of transcription factors and chromatin modifying proteins to the DSB site, thereby inducing a coordinated DNA damage response at the transcriptional level while repair takes place. Factors known to relocalize include the histone deacetylases SIRT1, SIRT6, HDAC1, and poly-ADP ribose polymerase, PARP1 (Dobbin et al., 2013; Mao et al., 2011; Oberdoerffer et al., 2008). After repair, the majority of the original chromatin structure is restored, but repeated triggering of the response disrupts youthful gene expression patterns and can de-silence retrotransposons that are held at bay by SIRT1 and SIRT6 (De Cecco et al., 2019; Oberdoerffer et al., 2008; Simon et al., 2019), analogous to erosion of the Waddington landscape or accumulation of epigenetic noise. Why the epigenetic clock is advanced by DSBs remains a mystery, but one possibility is that DSBs cause the relocalization of ten eleven translocation enzymes (Tets) or DNA methyltransferases (DNMTs) to DNA breaks, leading to specific changes in DNA methylation patterns over time (Field et al., 2018).

We cannot rule out that some of the effects we see in the ICE mice are due to cutting of the rDNA locus. Indeed, rDNA instability is a known cause of aging in yeast and the nucleolus size predicts lifespan (Sinclair et al., 1997; Tiku et al., 2017). Given that we saw no evidence of rDNA mutations, changes in rRNA levels or protein translation, and in light of the observation that the I-*Sce*I enzyme, which cuts at different genomic sites, generated a similar gene expression pattern (Yang et al., co-submitted manuscript), rDNA instability is unable to explain the ICE phenotype. Future work will include testing of CRISPR-based systems that do not induce mutations or cell cycle arrest (Chen et al., 2017; Kleinstiver et al., 2016) or those that direct DNA damage signaling proteins to a locus without activating a DSB signal (Soutoglou and Misteli, 2008).

The duality of decreased silencing and increased DNA repair was first detected in budding yeast, in which the derepression of silent mating-type genes increases DNA repair efficiency (Lee et al., 1999) but constitutive derepression causes a loss of cell identity and sterility, a hallmark of yeast aging (Smeal et al., 1996). This is a clear example of antagonistic pleiotropy, in which an adaptive process that promotes survival in young individuals disrupts homeostasis at an advanced age where the influence of natural selection falls to near zero.

Besides facilitating a new understanding of epigenetic change during aging, the ICE system may overcome a variety of other research challenges. Short-lived species such as rodents have proven to be poor models of many human age-related diseases. ICE mice, however, may solve this problem by accelerating the epigenetic age of the mice. Indeed, some of the phenotypes of the ICE mice, such as retinal degeneration, loss of vascularity and memory loss, closely resemble aging in humans. And by accelerating aging in specific tissues, it will be possible to test how much individual organ systems contribute to aging. By introducing the ICE system into human iPSCs, it should be possible to generate human tissue cultures and organoids that recapitulate human diseases of aging better than those that are currently available.

Because DSBs accelerate epigenetic clock and the aging process, it is possible the two processes are intimately linked. Whether or not it is possible to reverse the clock, recover lost epigenetic information, and restore the function of tissues remains to be seen but recent evidence indicates that it could be possible (Lu et al., 2019; Ocampo et al., 2016). If so, an understanding of why we age and potentially how to reverse it may be in reach.

## Supporting information

Supplemental Information_Hayano et al

## ACKNOWLEDGMENTS

This paper is dedicated to the memory of our colleague and co-author Norman S. Wolf. We thank to all members of the Sinclair laboratory for constructive comments. We thank Edward Schulak for financial support and encouragement, Andrea Di Francesco, Phu Huynh, Kristal Kalafut, Erin Wade and Rafael de Cabo (National Institute on Aging) for technical advice and Alex Banks (Beth Israel Deaconess Medical Center) for help with MRI. Supported by NIH/NIA (R01AG019719 and R37AG028730 to D.A.S), the Glenn Foundation for Medical Research (to D.A.S. and A.J.W.), Human Frontier Science Program (LT000680/2014-L to M.H.), JSPS KAKENHI (17K13228, 19K16619 and 19H05269 to M.H.), the Uehara Memorial Foundation (to M.H.), National Research Foundation of Korea (2012R1A6A3A03040476 to J.-H.Y.), NIH T32 (T32AG023480 to D.L.V.), NIA K99/00 (K99AG055683 to J.M.R.), NEI (RO1EY019703 to T.C.J.) and St Vincent de Paul Foundation (to B.R.K.).

## AUTHOR CONTRIBUTIONS

M.H., J.-H.Y. and D.A.S. designed, performed and analyzed a majority of the experiments. M.S.B., J.A.A., P.T.G. and D.L.V. helped write the manuscript. J.A.A. and M.S.B. conducted electron microscopy. J.M.R., J.A.A., G.C. and M.A.R. performed brain analyses and behavioral tests. Y.C.C., W.G., and X.Y. calculated epigenetic ages. P.T.G., D.L.V. and E.L.S. analyzed RNA-seq and WGS data. S.J.M. and A.E.K. assessed mouse frailty. Y.M., E.K.N., M.S.B. and G.F.M. performed skin IHC. L.Z. and R.M. helped metaphase spread. N.S.W., M.G.-K., T.C.J. and B.R.K. assessed eye phenotypes. H.W., J.G.S. and C.E.S. performed echocardiograms. J.A.K. and J.M.S. measured activation of repetitive sequences. A.D., S.T., N.G., A.-M.B., S.J.B., and L.S. conducted mouse phenotype analyses. K.T., C.M.P., and A.J.W. provided advice and assistance with experiments. P.O. and D.A.S. designed the I-*Ppo*I construct and generated the I-*Ppo*I mouse. L.A.R. initiated ICE mouse study and provide advice and assistance throughout. D.A.S. designed and supervised the study. J.-H.Y., M.H., and D.A.S. wrote the manuscript.

## METHOD DETAILS

### Mice and treatments

I-*Ppo*I^STOP^ knock-in mouse ES cells were generated the following way. Briefly, an estrogen receptor nuclear translocation domain (ER^T2^) tagged with HA at N-terminus and I-*Ppo*I were inserted into STOP-eGFP-ROSA26TV plasmid (addgene, plasmid #11739) together followed by IRES and EGFP sequence. HA-ER^T2^-I-*Ppo*I^STOP^ cassette was integrated at Rosa26 loci and the targeted C57BL6 ES cells were injected into C57BL/6 albino (cBRD/cBRD) blastocysts. After back-crossing I-*Ppo*I ^STOP/+^ chimeric mice with C57BL/6 mice, ICE mice were generated by crossing I-*Ppo*I ^STOP/+^ mice to Cre^ERT2/+^ mice harboring a single ER^T2^ fused to Cre recombinase that is induced whole body (Ruzankina et al., 2007). 4-6 month-old Cre and ICE mice were fed a modified AIN-93G purified rodent diet with 360 mg/kg Tamoxifen citrate for 3 weeks to carry out I-*Ppo*I induction. ER^T2^ containing three mutations selectively binds to 4-hydroxytamoxifen (4-OHT) but not estradiol. Cre-ER^T2^ protein is translocated into nucleus by tamoxifen treatment followed by removal of the STOP cassette located at upstream of I-*Ppo*I. In the presence of tamoxifen, Cre-ER^T2^ and HA-ER^T2^-I-*Ppo*I localize to the nucleus and induce DNA double strand breaks. Wild-type aged mice were obtained from the NIA aged rodent colonies and acclimatized at least for a month prior to experimentation. Mice were fed LabDiet 5053 diet and all animal care followed the guidelines of Animal Care and Use Committees (IACUCs) at Harvard Medical School.

### Cell culture

Cells were cultured as described in Yang et al. (co-submitted manuscript)

### Western blot analysis

Cell and tissue samples were lysed in RIPA buffer (50 mM Tris-HCl pH 7.4, 150 mM NaCl, 0.25% deoxycholic acid, 1% NP-40, 1 mM EDTA) containing a proteinase inhibitor cocktail (Sigma-Aldrich). An equal amount of lysate was incubated with sample buffer (0.05% Bromophenol blue, 2% sodium dodecyl sulfate, 50 mM Tris-Cl pH 6.8, 5% β-mercaptoethanol) at 95°C for 5 min then separated on an SDS-PAGE gradient gel, transferred to a membrane using transfer buffer (25 mM Tris-HCl pH 8.3, 190 mM glycine 20% methanol), blocked with TBSTM (Tris-buffered saline, 0.1% Tween 20, with 5% skim milk), probed with primary and secondary antibodies and developed using ECL Western Blotting Detection Reagent (Sigma-Aldrich).

### Comet assay

A comet assay was performed using the OxiSelect Comet Assay Kit (Cell Biolabs). In brief, 10^4^ cells were resuspended in 100 µl liquid agarose and 75 µl was immediately dropped onto the comet slides, then incubated at 4°C for 15 min to solidify the agarose. Slides were submerged in pre-chilled lysis buffer at 4°C for 45 min, transferred to alkaline solution (300 mM NaOH, pH >13, 1 mM EDTA) and stored at 4°C for 60 min in the dark. Electrophoresis was performed with cold alkaline solution, then slides were washed with water and 70% ethanol. After air drying the agarose, DNA was stained with Vista Green and Comet olive tail moments (Tail DNA% × Tail Moment Length) were measured using CASPLab (http://casplab.com/) software.

### Southern blotting

Genomic DNA samples were prepared using EZNATissue DNA Kit (Omega Bio-tek). DNA (3 µg) was run in 0.8% agarose gel, DNA was depurinated in 0.25 N HCl, denatured in 0.4 N NaOH, and washed with 20X SSC. DNA was transferred to a nylon membrane in 0.4 N NaOH using a TurboBlotter (Whatman), washed with 2X SSC, crosslinked by UV then incubated in pre-hybridization solution (6X SSC, 5X Denhardt’s solution, 1X SSD,0.0625 M Tris-HCl pH7 .5, 75 µ/ml salmon sperm DNA) at 65°C for 3 h with rotation. DNA probes were generated using target-specific PCR with dCTP [α-32P]. Radioactive DNA probes were added to fresh pre-hybridization solution and incubated with the membrane overnight with rotation. The membrane was washed with 2X SSC, 2X SSC containing 1% SDS and 0.1X SSC and exposed to X-ray film at −80°C.

### Surveyor assay

I-*Ppo*I target regions were amplified from genomic DNA isolated from either Cre or ICE cells by PCR using flanking primer sets. Hetero- or homo-duplexes were hybridized in thermocycler and hybridized DNA (200 ng) was treated with SURVEYOR nuclease S (Transgenomic) at 42°C for 60 min. Nuclease reactions were stopped and digestion was analyzed by agarose gel electrophoresis or a Bioanalyzer (Agilent).

### Metabolic labeling of MEFs

MEFs were washed twice with pulse-labeling medium (Met-Cys-free DMEM containing 10% dialyzed serum) and incubated in pulse-labeling medium for 1 h to deplete intracellular Methionine. Pulse-labeling medium with 0.2 mCi/ml methionine [35S] was added to cells and incubated for 1h. Cells were lysed and ^35^S-methionine incorporation was determined by TCA precipitation and scintillation counting.

### Metaphase spread and telomere FISH

Cells were treated with Colcemid (10 ug/ml, Gibco) for 4 h. Trypsinized cells were washed with PBS and mixed in prewarmed 4 ml of 0.075M KCl by vortexing. After 15 min incubation, fixative solution (methanol:acetic acid = 3:1) was added. Cells were dropped in steam-moisturized slide glasses and dried on hot plate. Cells were fixed in 4% formaldehyde for 2 min, then digested in pepsin for 10 min. Another round of fixing was performed and cells were dehydrated in 70%, 90%, 100% EtOH for 5 min each. 0.5 µg/ml telomere PNA FISH probe (PNA BIO) was prepared and incubated at 80°C for 3 min. The probe was added to the slides and incubated at RT for 2 h. Slides were washed, dehydrated and dried, and nuclei were stained with antifade mounting medium containing DAPI (Vector Laboratories).

### Quantification of DSBs

DNA double strand breaks (DSB) generated by I-*Ppo*I were detected as described previously (Chailleux et al., 2014). Briefly, tissue was homogenized in phenol and genomic DNA was purified with chloroform, ethanol and RNase. Genomic DNA carrying I-*Ppo*I specific DSBs was subjected to ligation-mediated purification using biotin-conjugated adaptor nucleotides with 5‘-AATT-3’ overhangs that bind to the DSB site generated by I-*Ppo*I. Adaptor sequences were as follows: dRbiot-BglII-IPpoI F 5‘-CCCTATAGTGAGTCGTATTAGATCTGCGTTAA-3’, dRbiot-BglII-IPpoI R 5‘-CGCAGATCTTAATACGACTCACTATAGGG-3’. The biotinylated fragment was digested using *Eco*RI for 3 h at 37°C followed by purification with streptavidin magnetic beads (Dynabeads M-280 Streptavidin, Invitrogen) in binding buffer (20 mM Tris-HCl pH 8.0, 0.1% SDS, 1% Triton X-100, 2mM EDTA, 150 mM NaCl). After 4 h at 4°C, beads were washed five times with washing buffer (50 mM Tris-HCl pH 8.0, 0.1% SDS, 150 mM NaCl) and once with TE buffer. Cut DNA was eluted by digesting the adaptor with *Bgl*II at 37°C overnight. DNA was purified using glycogen, sodium acetate and ethanol. DNA primers were: 5+11 F 5‘-ACTTAGAACTGGCGCTGAC-3’, 5+11 R 5‘-CTGGCCTGGAACTCAGAAAT-3’, 28S F CCCACTGTCCCTACCTACTATC, 28S R AGCTCAACAGGGTCTTCTTTC.

### Quantification of protein synthesis

Protein synthesis was quantified as published (Garlick et al., 1980; Hofmann et al., 2015). L-^3^H-phenylalanine (1 mCi/mL) was combined with unlabeled phenylalanine (135 mM) to create 100 mCi/ml. After adjusting the solution to pH 7.1 with NaOH, the labeling solution was injected via the lateral tail vein at 1 ml/100 g bodyweight under anesthesia with ketamine (75 mg/kg) and xylazine (10 mg/kg).

### Indirect Calorimetry

Food consumption, ambulatory activity, oxygen consumption (VO_2_), carbon dioxide production (VCO_2_) and respiration exchange ratio (RER) were measured using Columbus Instruments CLAMS. Mice were housed in metabolic cages for 3 days prior to collecting data and body composition was determined by EchoMRI 3-in-1.

### MMQPCR

Monochrome multiplex quantitative PCR was performed as described previously (Cawthon, 2009). Briefly, a PCR reaction containing 20 ng of genomic DNA was prepared with SYBR Green system (Applied Biosystems). The PCR program was set up as Step 1: 95°C; 15 min, Step 2; 2 cycles of 94°C for 15 sec and 49°C for 15 sec, Step 3: 32 cycles of 94°C for 15 sec, 62°C for 15 sec, 74°C for 15 sec with signal acquisition for telomere or 28S amplification, 84°C for 10 sec, 88°C for 15 sec with signal acquisition for Hbbt1 amplification. Primers are listed in Table 1.

**Table 1.**

| REAGENT or RESOURCE | SOURCE | IDENTIFIER |
| --- | --- | --- |
| **Antibodies** |  |  |
| Rabbit polyclonal anti-γH2AX | Abcam | Cat# ab2893 |
| Rabbit polyclonal anti-γH2AX | Cell Signaling Technology | Cat #2577 |
| Mouse monoclonal anti-γH2AX | Novus | Cat# NBP1-19255 |
| Rabbit polyclonal anti-H2AX | Abcam | Cat# ab11175 |
| Rabbit polyclonal anti-p53p | Cell Signaling Technology | Cat# 9284 |
| Mouse monoclonal anti-P53 | Cell Signaling Technology | Cat# 2524 |
| Mouse monoclonal anti-ATMp | Cell Signaling Technology | Cat# 4526 |
| Rabbit polyclonal anti-p16 | Santa Cruz Biotechnology | Cat# sc-1207 |
| Rabbit polyclonal anti-PARP1 | Cell Signaling Technology | Cat# 9542 |
| Rat monoclonal anti-HA-Peroxidase | Roche | Cat# 12013819001 |
| Rat monoclonal anti-CD31 (RM0032-1D12) | Abcam | Cat# ab56299 |
| Rabbit polyclonal anti-Laminin | Sigma-Aldrich | Cat# L9393 |
| Rabbit polyclonal anti-Iba1 | Wako | Cat# 01919741 |
| Rabbit polyclonal anti-GFAP | Abcam | Cat# A85419 |
| Alexa Fluor 594 Goat anti-rabbit (H+L) | Life Technologies | Cat# A11012 |
| Alexa Fluor 488 Goat anti-rabbit IgG (H+L) | Life Technologies | Cat# R37116 |
| **Chemicals, Peptides, and Recombinant Proteins** |  |  |
| Tamoxifen citrate salt | Sigma-Aldrich | Cat# T9262 |
| (Z)-4-Hydroxytamoxifen (4-OHT) | SIGMA | Cat# H7904 |
| Etoposide | CALBIOCHEM | Cat# 341205 |
| Camptothecin | CALBIOCHEM | Cat# 208925 |
| Paraquat | SIGMA | Cat# 36541 |
| Hydrogen peroxide | SIGMA | Cat# 216763 |
| Rodent Chow Diet | LabDiet | Cat# 5053 |
| LightCycler 480 SYBR Green I Master | Roche | Cat# 4707516001 |
| Dynabeads® Protein A for Immunoprecipitation | Life Technologies | Cat# 10001D |
| Dynabeads® Protein G for Immunoprecipitation | Life Technologies | Cat# 10003D |
| I-Ppol | Promega | Cat# R7031 |
| Dynabeads® M-280 Streptavidin | Life Technologies | Cat# 11205D |
| UltraPure™ Buffer-Saturated Phenol | Thermo Fisher | Cat# 15513039 |
| Phenylalanine, L-[2,3,4,5,6-3H] | PerkinElmer | Cat# NET1122001MC |
| Hematoxylin solution modified acc. to Gill II | Millipore | Cat# 1051750500 |
| Eosin Y-solution 0.5% alcoholic | Millipore | Cat# 1024390500 |
| Fluoroshield mounting medium with DAPI | Sigma-Aldrich | Cat# F6057 |
| Colcemid | Thermo Fisher Scientific | Cat# 15212012 |
| Pepsine | SIGMA | Cat# P7000 |
| **Oligonucleotides** |  |  |
| Telomere PNA FISH probe | PNA BIO | Cat# F1006 |
| Experimental Models:<br>Organisms/Strains |  |  |
| C57BL/6 mouse | NIA (USA) | N/A |
| C57BL/6 HA-ERT2-I-*Ppol* mouse | This paper | N/A |
| C57BL/6 Cre-ERT2 mouse | Ruzankina et al., 2007 | N/A |
| **Software and Algorithms** |  |  |
| GraphPad Prism v8 | GraphPad Software | https://www.graphpad.com/scientific-software/prism/ |
| Primer-BLAST | NIH | https://www.ncbi.nlm.nih.gov/tools/primer-blast/ |
| Cellprofiler | Broad Institute | http://cellprofiler.org/ |

### Frailty index assessment

The Frailty Index (FI) was scored as described previously (Whitehead et al., 2013). Briefly 31 health-related deficits were assessed for each mouse. A mouse was weighed and body surface temperature was measured three times with an infrared thermometer (La Crosse Technology). Body weight and temperature were scored based on their deviation from mean weight and temperature of young mice (Whitehead et al 2014). Twenty-nine other items across the integument, physical/musculoskeletal, oscular/nasal, digestive/urogenital and respiratory systems were scored as 0, 0.5 and 1 based on the severity of the deficit. Total score across the items was divided by the number of items measured to give a frailty index score between 0 and 1.

### Lens opacity scoring

Lens opacity scoring was previously described (Wolf et al., 2008). Mice were held without anesthesia and assessed in a dark room using a SL-14 Kowa hand-held slit lamp (Kowa, Tokyo, Japan).

### PET-CT

Mice were anesthetized with 2% isoflurane gas in oxygen and injected with ∼200 µCi F-18 labeled flourodeoxyglucose (FDG) via tail vein injection. After 45 minutes, mice were imaged on a Siemens Inveon small animal imaging scanner for positron emission tomography (PET) and computed tomography (CT) imaging under isoflurane. CT imaging was performed over 360 projections with a 80 kVp 500 µA x-ray tube and reconstructed using a modified feldkamp cone beam reconstruction algorithm (COBRA Exxim Inc., Pleasanton, CA) with 425 ms exposure time/projection during which Isovue-360 (Bracco Diagnostic Inc, Monroe Township, NJ) was pumped into the mouse via tail vein at a rate of 20 µl per minute. PET scans were reconstructed with ordered subset expectation maximization with maximum a posterior reconstruction algorithm with 2 OSEM iterations and 18 MAP iterations.

### Magnetic Resonance Imaging

Mice were anesthetized with 2% isoflurane gas in oxygen and placed in a 4.7 Tesla Bruker Pharamscan magnetic resonance imager. Rare T1 (TE: 13.4 ms, TR: 900 ms, Rare factor: 4, Matrix: 256 × 256 × 24, Voxel size: 0.215 × 0.156 × 1 mm) and a Rare T2 (TE: 18.26 ms, TR: 2000 ms, Rare factor: 8, Matrix: 256 × 256 × 24, Voxel size: 0.215 × 0.156 × 1 mm) scans of the lower thoracic cavity, abdomen and lower extremities were performed.

### Micro CT scanning

Femurs were isolated and placed in 70% ethanol. Micro-CT was performed by using SCANCO Medical μ-CT35 at the core facility at the Harvard School of Dental Medicine (Idelevich et al., 2018).

### Quantification of optic nerve axons

To quantify axons, optic nerves were dissected and fixed in Karnovsky’s reagent (50% in phosphate buffer) overnight. Semi-thin cross-sections of the nerve were taken at 1.0 mm posterior to the globe and stained with 1% p-phenylenediamine (PPD) for evaluation by light microscopy. Six to eight non-overlapping photomicrographs were taken at 60x magnification covering the entire area of the optic nerve cross-section. Using ImageJ software, a 100 × 100 μM square was placed on each 60x image and all axons within the square (0.01 mm^2^) were counted using the threshold and analyze particles function in image J. The average axon counts in 6-8 images was used to calculate the axon density/mm^2^ of optic nerve. Scorers were blinded to experimental groups.

### Immunohistochemistry for mouse skin

Dorsal skin samples were fixed with 4% paraformaldehyde/PBS and kept on ice for 2 h. The fixed skin samples were embedded in OCT (Sakura Finetek) and snap frozen in liquid nitrogen for histology. After washing in PBS, nonspecific staining was blocked with PBS containing 3% skim milk and 0.1% Triton-X for 30 min. Sections were incubated with primary antibodies at 4°C overnight: rat anti-mouse CD117 (BD Pharmingen) and rabbit anti-human KRT5 (COVANCE). Secondary antibodies were conjugated with Alexa Fluor 488 or 594 (Invitrogen). Nuclei were counterstained with 4’,6-diamidino-2-phenylindole (DAPI) and images were obtained using FV1000 confocal microscope (Olympus). >100 hair follicles per mouse (n=8) were analyzed for the presence of KIT+ melanocytes in the bulge.

### Quantification of subepidermal thickness

Site-matched skin tissue was fixed in formalin, embedded in paraffin, and 5 µm sections were cut and stained with hematoxylin and eosin. Representative regions of the subcutaneous layer were measured from the limits of the dermis to the panniculus carnosus (‘subepidermis’) with the assistance of an ocular micrometer. Care was taken to ensure that tissue was embedded perpendicularly and the subdermal thickness determination was not artificially enhanced due to tangential sectioning. Because the subepidermal layer reached maximum thickness in control Cre mice at 17-18 months, this timepoint was selected for comparisons with the ICE mice. A minimum of 10 randomly selected thickness determinations were generated for each tissue section.

### Brain immunohistochemistry

For GFAP and Iba1 staining, the tissues were incubated overnight in paraformaldehyde (4% v/v). Fixed brains were embedded in paraffin and 6 µm sections were generated using a manual rotary microtome (Leica). After deparaffinization and re-hydration of tissue slides, an antigen revealing step was performed by using antigen unmasking solution (Vector). Sections were blocked in PBS with 5% BSA and 0.3% Triton-X at 4°C for 1 hr and incubated with primary antibodies in PBS with 2% BSA and 0.1% Triton-X at 4°C overnight with Rabbit anti-GFAP antibody (Abcam, ab7260), Rabbit anti-Iba1 antibody (Funakoshi, GTX100042). Secondary antibodies conjugated with Alexa Fluor 488 or 594 (Invitrogen) were used followed by DPAI staining. To localize I*-Ppo*I expression and DNA damage, mice were perfused transcardially and brains were post-fixed overnight with 4% paraformaldehyde/PBS, then cleared by 30% sucrose solution. Brains were embedded in OCT (Sakura Finetek) and 40 µm sections were collected using a cryostat (Leica). Sections were blocked in horse serum/TBS-Triton-X for 30 min at RT, and then incubated with primary antibodies overnight at 4°C with goat anti-GFP (Abcam) and rabbit anti-γ-H2AX (Cell Signaling). Secondary antibodies were conjugated with Alexa Flour 488 and 647 (Jackson ImmunoResearch) and co-stained with DAPI.

### ATP and mtDNA measurement

Snap frozen tissue was briefly washed with PBS and 3 ml Tris-HCl TE saturated phenol per 100 mg was added to the tissue followed by homogenizing with a tissue homogenizer (Omni TH, Omni). After centrifugation, cell lysates were added to an equal amount of TE saturated phenol, chloroform and water were added to the same tube. After centrifugation, the supernatant was used for ATP and mtDNA measurement. ATP was measured using an ATP kit (ThermoFisher Scientific) and normalized to tissue weight. Genomic DNA and mtDNA were purified with 2.5-fold ethanol and glycogen. Primers for 18S ribosomal and CytB were used to calculate the ratio of mtDNA to genomic DNA. Primers were: mouse 18S, 5‘-TGTGTTAGGGGACTGGTGGACA-3’ (forward) and 5‘-CATCACCCACTTACCCCCAAAA-3’ (reverse), mouse Cytb, 5‘-CCCTAGCAATCGTTCACCTC-3’ and 5‘-TGGGTCTCCTAGTATGTCTGG-3’ (reverse).

### Open field test

Mice were placed in the center of a chamber (48.5 cm × 48.5 cm) enclosed by a curtain. Novel environment exploration was assessed by allowing mice to explore for 5 min. Exploratory activity at the periphery and total activity were recorded using a video tracking system and analyzed by TopScanLite version 2.

### Contextual fear conditioning test

Contextual fear conditioning was assessed using a TSE system. On day 1, mice were placed into an experimental box (52 cm × 52 cm × 65 cm) and allowed to explore freely for 180 s followed by 0.5 mA electric shock for 1 s. One more 0.5 mA shock for 1 s was given after 30 s and immediate freezing was measured every 10 s by a visual count, after which mice were returned to their home cage. Contextual freezing without a tone was assessed for 180s, 24 hours after the shock, counting freezing every 10 sec.

### Barnes maze test

The maze consisted of a circular and white platform (90 cm in diameter) with 20 × 5 cm diameter holes arranged around the edge of the platform, elevated 82 cm above the floor. For visual cues, the platform was surrounded by four pictures with different colors and shapes. A mouse was placed in the center of maze and then, the mouse was guided to a small chamber termed a target hole at adaptation period. After 2 min in the target hole, the mouse was returned to the cage. During the spatial acquisition period, the mouse was allowed to explore the target hole for 3 min. If the mouse entered the target hole or it passed 3 min, the mouse was left for 1 min in the hole. The trial was repeated 3 times/day for 5 d. A probe trial was performed to test long-term memory 7 days later by covering the target hole with a lid. The mouse was allowed to explore the position of target hole for 90 s and the number of pokes in each hole was measured using TopScanLite version 2.

### Grip strength test, treadmill test and lactate measurement

To measure muscular strength, a mouse was held by the tail and allowed to grip a mesh grip with the front paws (BIO-G53, BIOSEB) then pulled backward until grip was released. After a 10 min break, the experiment was repeated. Maximum exercise endurance was assessed with a treadmill system (TSE). Mice were trained for 3 d prior to recording the performance to familiarize the mice to the equipment. An electrical stimulation grid was adjusted as 1 mA and slope was set at 15 degrees. The first day of the training, mice walked on the treadmill at 10 m/min speed for 10 min, with a 10 min break, then walked at 10 m/min speed for 10 min. On the second and third day, the initial two steps were the same as first day, then walking was started at 10 m/min and the speed was increased by 1 m/min every minute to a maximum speed of 20 m/min. On day 4, maximum exercise endurance was measured. Six mice were placed on the treadmill and the belt speed was started at 5 m/min for 5 min to allow the mice warm up. The speed was increased by 1 m/min up to 20 m/min. After running for 5 min, the speed was increased from 20 m/min to 21 m/min for 10 min. Mice were then forced to run at 22 m/min until they remained on the electrical stimulation grid for 10 seconds. Details are available upon request. The tail blood at pre-exercising and post-exercising was taken and serum lactate level were measured with a lactate meter (Nova Biomedical).

### Ambulatory activity

Animals were maintained in specific-pathogen-free (SPF) facility and single-housed in instrumented individually ventilated cages (IVC) (Digital Smart House, Vium, San Mateo, CA, and Innovive, San Diego, CA) containing corncob bedding with access to Innowheel and Innodome (Innovive, San Diego, CA), Bed-r’Nest (Andersons Lab Bedding, Maumee, OH), and foraging mix (Veggie Relish, LabDiet). Animals had unrestricted access to food (Pico Rodent Diet 5053, Lab Diet, St. Louis, MO) and acidified, sterile water (Innovive, San Diego, CA). Vium Digital Smart Houses slotted in Vium’s proprietary rack system were outfitted with sensors and a high-definition (HD) camera that enables continuous, 24/7 monitoring of animals and streams data to a secure cloud-based infrastructure. As described elsewhere (Lim et al., 2019; Lim et al., 2017), video is processed using computer vision algorithms to produce a digital history of motion (mm/sec). Motion (mm/s) was averaged across 1 hr bins to produce 1 hr averages. All 1 hr averages from 6am to 7am across 55 days were averaged and repeated for each hour of the day.

### Whole genome sequencing (WGS)

Genomic DNA was isolated from snap frozen tissues using a DNeasy blood & tissue kit and fragmented with a Covaris ultrasonicator at 500 bp peak. TruSeq DNA PCR-free library preparation kits were used to add DNA adaptors to dsDNA following manufacturer’s instructions (Illumina). Deep whole genome sequencing (50X) on an Illumina Hiseq X10 platform was performed at BGI (China).

### Treadmill Gait Analysis

Gait patterns were measured using forced walking on a treadmill (Columbus Instruments; Columbus, OH). A high-speed digital video camera recorded images of the ventral side of the mouse through a transparent treadmill belt reflected off a mirror. Mice for approximately 24 sec at speeds of 13, 19, and 25 cm/s. TreadScan® software (CleverSys, Inc, Reston, VA) identified each individual paw of the mouse in each frame as it walked on the treadmill and measures of stance and swing duration, among other measures, were assessed.

### COX and capillary density staining

Freshly isolated quadriceps and gastrocnemius muscles were mounted in OCT (Tissue-Tek), placed in an isopentane bath, and slowly cooled in liquid nitrogen. Transverse sections (20 mm) were sectioned on a cryostat (Leica). Sections were fixed in pre-cooled acetone (−20°C) for 10 min, washed with PBS, then blocked with BlockAid (Invitrogen) for 1 h at RT, and then incubated with CD31 (ab56299, Abcam), Laminin (L9393, Sigma) antibodies diluted in blocking buffer overnight at 4°C. Slides were washed with PBST, then incubated with anti-rat Alexa Fluor 488-conjugated (Life Technologies) and anti-rabbit Alexa Fluor 594-conjugated (Life Technologies) diluted to 1:500 in blocking buffer for 2 h at RT. Slides were washed again with PBST and mounted with Fluoroshield with DAPI mounting medium (Sigma). Images were acquired using a confocal fluorescence microscope (Nikon A1). COX staining was performed according to a protocol (Ross, 2011). Briefly, 20 μm cryostat sections was dried at room temperature for 1 hr and media containing 1X DAB, 100 μM cytochrome c, 2 μg/ml bovine catalase was added to sections and slides were incubated at 37°C for 40 min. Quantification of capillary number and density were performed using ImageJ.

### Electron microscopy

Mice at 15 months of age were anesthetized with isoflurane and sacrificed by cervical dislocation or decapitation, in accordance with available ethical permits. Muscle was collected and fixed in electron microscopy fixative (consisting of 3% glutaraldehyde, 2.5% paraformaldehyde, 2 mM calcium chloride, 2% sucrose in 0.1 M cacodylate buffer) and tissue was processed as previously reported (Le Couteur et al., 2001). Two blocks from different parts of the muscle were used and from each section 10 images were taken at 5000X on a Jeol 1210 transmission microscope and photographed using a Gatan US 4000MP digital camera.

Mitochondrial network, size and number were quantified blindly using FUJI ImageJ.

### Quantitative real-time PCR for transcription of repetitive elements

Total RNA was isolated from 30-50 mg of tissue using Trizol reagent (ThermoFisher) according to the manufacturer’s instructions. Prior to the synthesis of cDNA, total RNA was digested with 27.2 Kunitz units of RNase-free DNase (Qiagen) for 45 min at room temperature and further cleaned up on RNeasy columns (Qiagen) (De Cecco et al., 2013). The effectiveness of the digestion was assessed using controls that omitted reverse transcriptase (RT). Digestion with DNase was repeated until the control lacking RT was negative for γ-satellite sequences. RNA integrity was determined using an Agilent Bioanalyzer 2100 and an RNA-nano chip. Total RNA (1 μg) of was transcribed into cDNA in 50 μl reactions using the TaqMan Gold RT-PCR kit (Applied Biosystems) and random hexamers, according to the manufacturer’s protocol. This reaction (1.0 μl) was used in subsequent qPCR reactions, performed using the SYBR Green system (Applied Biosystems) on the ViiA 7 Real Time System (Applied Biosystems), according to the manufacturer’s specifications. Primers were used at a final concentration of 300 nM. Tissue from 6 individual animals was analyzed in triplicate. Statistical analysis was determined using Student’s t-test and SigmaPlot 12.5 (Systat Software).

### Design of PCR primers for repetitive elements

All primers used in this study are listed in Table S1. For expression analysis of LINE-1, MusD and pericentromeric γ-satellite sequences (MSAT) we used primers described by Changolkar et al. (Changolkar et al., 2008). Primers for the SINE elements B1 and B2 were designed using the consensus sequence from Repbase (Genetic Information Research Institute, www.girinst.org/repbase/index.html) and Primer-Blast software (www.ncbi.nlm.nih.gov/tools/primer-blast/). Primers against GAPDH and β-actin, used as normalization controls, were designed with Primer-Blast using NCBI reference sequences NC_000072.6 and NM_007393.3, respectively. Primer sequences were analyzed using the UCSC genome browser *in silico* PCR tool (genome.ucsc.edu/cgi-bin/hgPcr) to determine the number of genomic elements that contribute to the amplification products (De Cecco et al., 2013). All primers were tested with serial dilutions of cDNA to ensure they amplified their target sequences quantitatively.

### Muscle RNA-seq analysis

Paired-end reads from gastrocnemius muscle RNA-Seq were mapped to the UCSC mm10 genome build using HISAT2 version 2.1.0 (Kim et al., 2015). The featureCounts function from the Rsubread package (Rsubread 1.32.2) was used to collect read counts for genes. DESeq2 (DESeq2 1.22.2) was applied for differential expression analysis to all genes with rowSums >= 10.

To compare gene expression in gastrocnemius muscles of ICE, Cre, and WT, a table of normalized read counts was exported from a combined DESeq dataset with all replicates and conditions. The 200 genes with the smallest adjusted p-value for differential expression between Cre and ICE were selected and ordered by the log2-fold-change difference between Cre and Ice. The heatmap.2 (gplots 3.0.1) R function was used to produce a plot of Z-score values for each gene.

### Epigenetic clock (DNAme age)

Tissue samples were immediately preserved in DNA/RNA Shield™ (Zymo Research; Cat. No. R1100-50) and genomic DNA were purified using Quick-DNA Plus Kit (Zymo Research; Cat. No. D4068) according to manufacturer’s instructions. Sample library preparation and data analyses were performed by Zymo Research, CA. Briefly, genomic DNA (200 ng) was bisulfite-converted using EZ DNA Methylation-Lightning™ Kit (Zymo Research; Cat. No. D5030).

Bisulfite-converted DNA libraries for targeted bisulfite sequencing platform, called SWARM^®^ (Simplified Whole-panel Amplification Reaction Method) were prepared according the to the manufacturer’s instructions, then sequenced on a HiSeq 1500 sequencer at >1,000X coverage. Sequence reads were identified using Illumina basecalling software and aligned to the reference genome using Bismark (http://www.bioinformatics.babraham.ac.uk/projects/bismark/), an aligner optimized for bisulfite sequence data and methylation calling. The methylation level of each sampled cytosine was estimated as the number of reads reporting a C, divided by the total number of reads reporting a C or T. DNA methylation levels of >500 age-related CpG loci were used for age prediction using epigenetic age algorithms.

## Supplemental Information

Table S1. Primers used in this study, Related to Figures 1, 2 and 5

Table S2. Differential genes from muscle RNA-seq, Related to Figure 7

Table S3. DNA methylation values of epigenetic clock CpGs, Related to Figure 7

---

**Figure S1.**
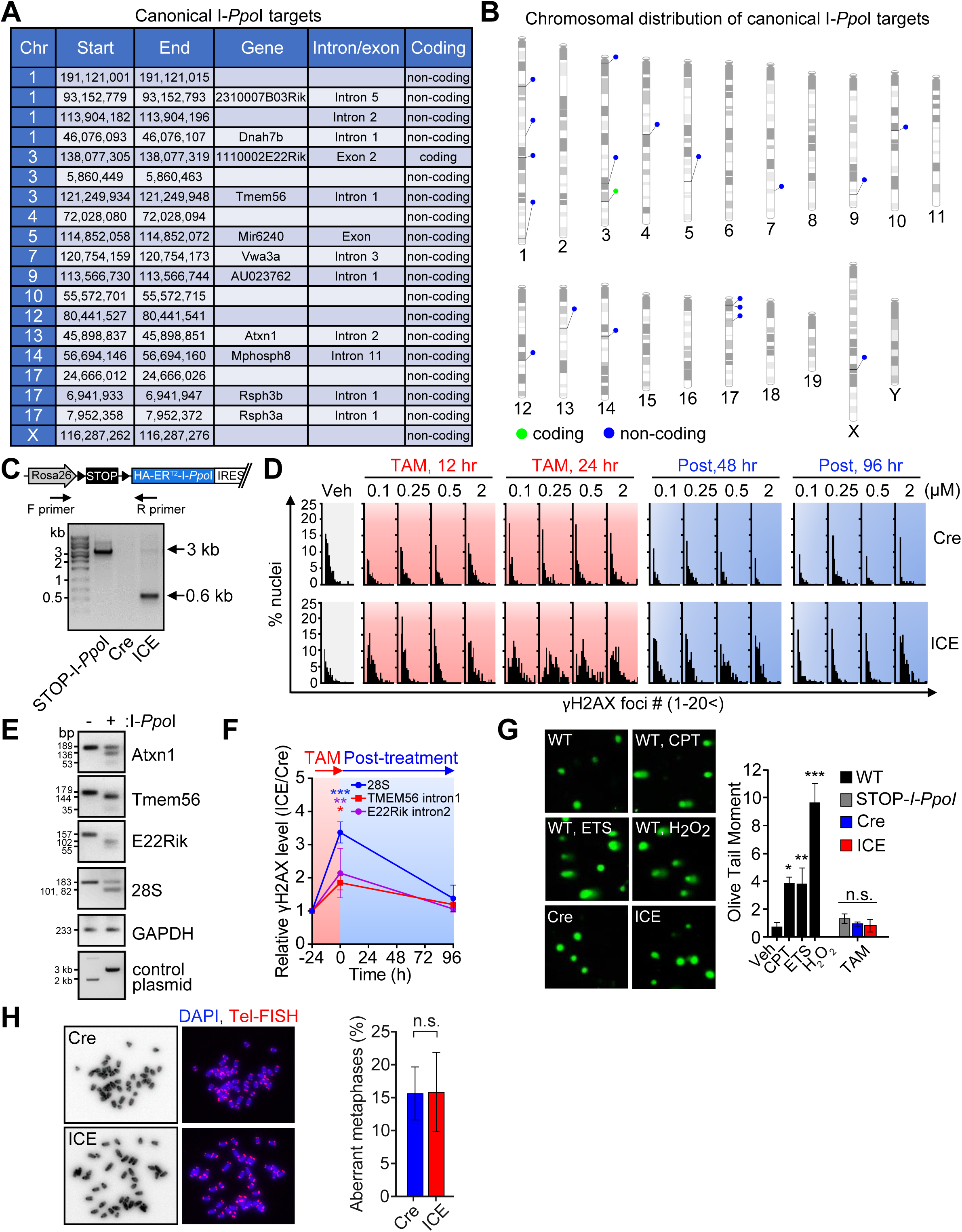
A Cell-Based ICE System Does Not Induce Genomic Instability, Related to Figure 1. (A) Canonical I-*Ppo*I targets in the mouse genome. (B) Genomic distribution of canonical I-*Ppo*I targets. (C) PCR analysis of STOP cassette excision during 4-OHT treatment. (D) Quantification of γH2AX foci induction during 4-OHT treatment and post-treatment. (E) *In vitro* cutting of PCR-amplified I-*Ppo*I targets using purified I-*Ppo*I. (F) Chromatin immunoprecipitation (ChIP) analysis of γH2AX deposition at I-*Ppo*I target sites. One-way ANOVA-Bonferroni. (G) Single cell electrophoresis of MEFs after treatment with camptothecin (CPT, 25 µg/ml), etoposide (ETS, 10 µM), hydrogen peroxide (H_2_O_2_, 100 µM), or I-*Ppo*I induction (4-OHT, 1 µM). One-way ANOVA-Bonferroni. (I) Percent of cells with aberrant metaphase chromosomes. Two-tailed Student’s *t* test. Data are mean (n=2-3) ± SEM. n.s.: p > 0.05; *p < 0.05; **p < 0.01; ***p< 0.001.

**Figure S2.**
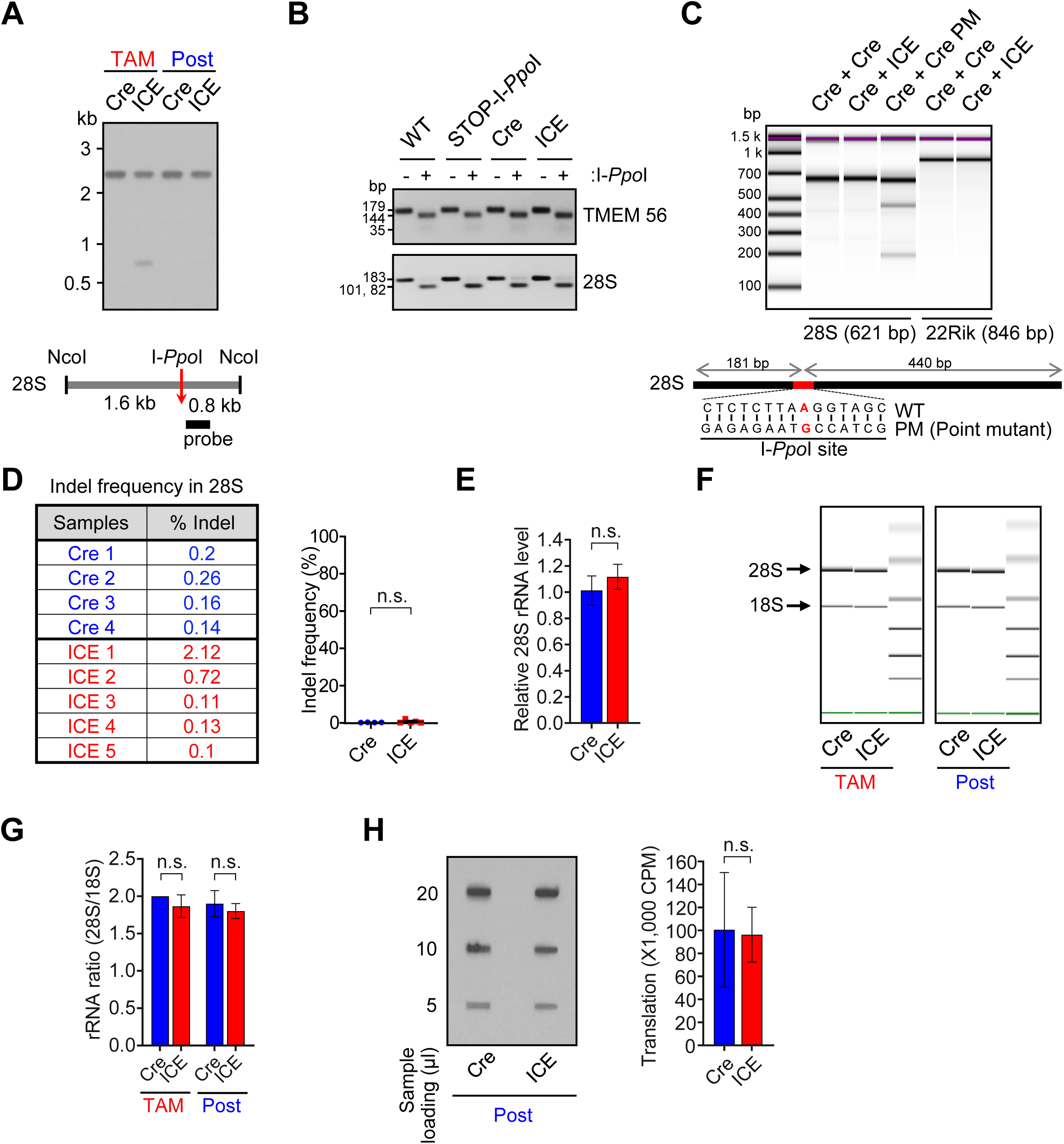
A Cell-Based ICE System Does Not Induce Mutations in rDNA, Related to Figure 1. (A) Southern blot of 28S rDNA during 4-OHT treatment and post-treatment. (B) *In vitro* cutting of I-*Ppo*I targets PCR-amplified from genomic DNA from post-treated ICE cells. (C) Surveyor nuclease assay of I-*Ppo*I targets in post-treated ICE cells. A PCR fragment with a point mutation (Cre PM) in the I-*Ppo*I site served as a positive control. (D) Mutation frequency of 28S rDNA in in post-treated ICE cells. Two-tailed Student’s *t* test. (E) 28S rRNA level in post-treated ICE cells. Two-tailed Student’s *t* test. (G) Bioanalyzer tracks of 28S and 18S rRNA in post-treated ICE cells. (G) 28S:18S rRNA ratio. Two-tailed Student’s *t* test. (H) Protein translation in post-treated ICE cells assessed by metabolic ^35^S-labelling. Two-tailed Student’s *t* test. Data are mean (n≥3) ± SD. n.s.: p > 0.05.

**Figure S3.**
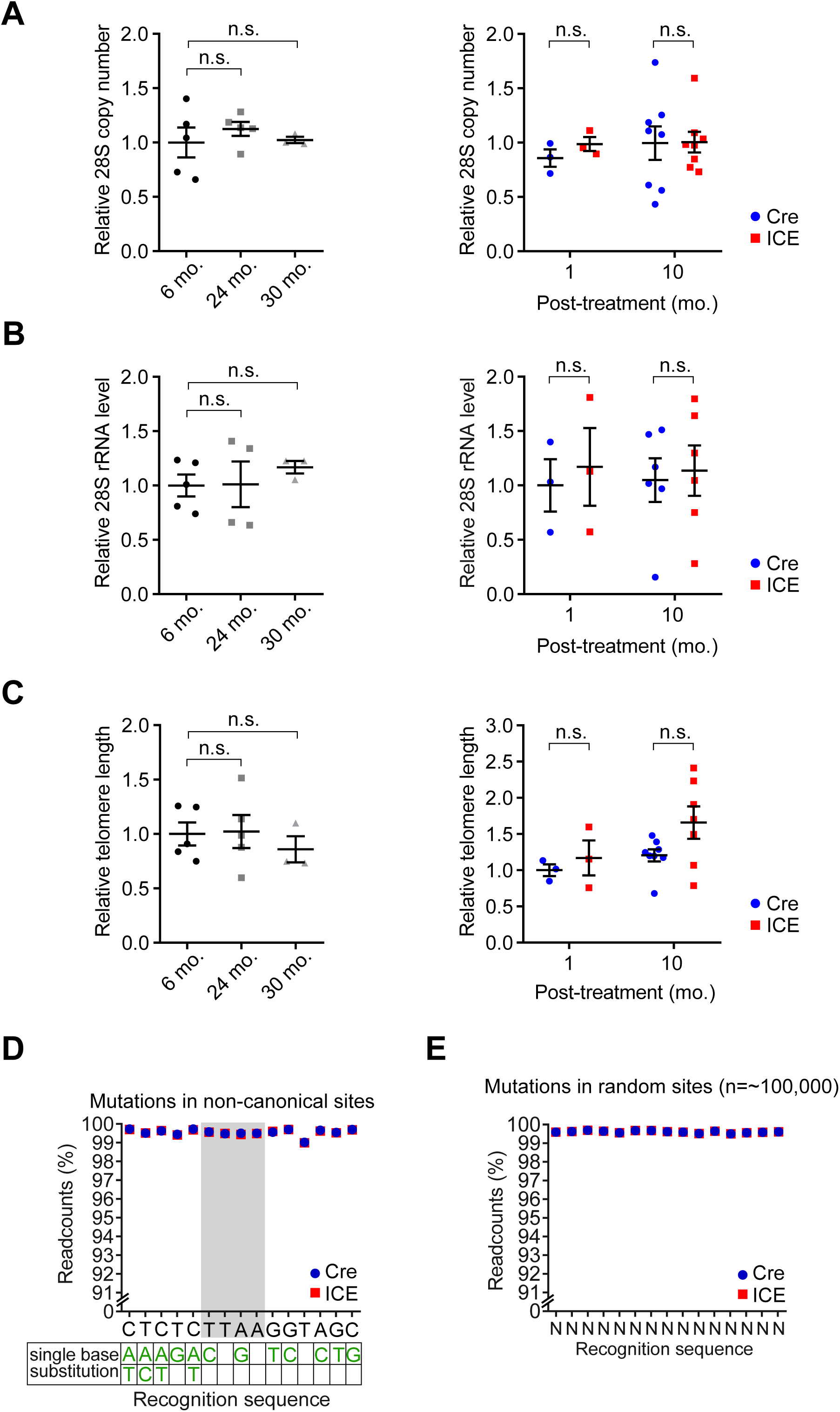
No Change in 28S rDNA or Telomere Lengths in Post-Treated ICE Muscle, Related to Figure 2. (A) 28S rDNA copy number assessed by monochrome multiplex quantitative PCR (MMQPCR). Two-way ANOVA-Bonferroni. (B) 28S rRNA levels. Two-way ANOVA-Bonferroni. (C) Telomere length assessed by monochrome multiplex quantitative PCR (MMQPCR). Two-way ANOVA-Bonferroni. (D and E) Percent non-mutated sequences in I-*Ppo*I non-canonical sites or ∼100,000 random sites by deep sequencing (>50x) in post-1-month ICE muscle. Data are mean ± SEM. n.s.: p > 0.05.

**Figure S4.**
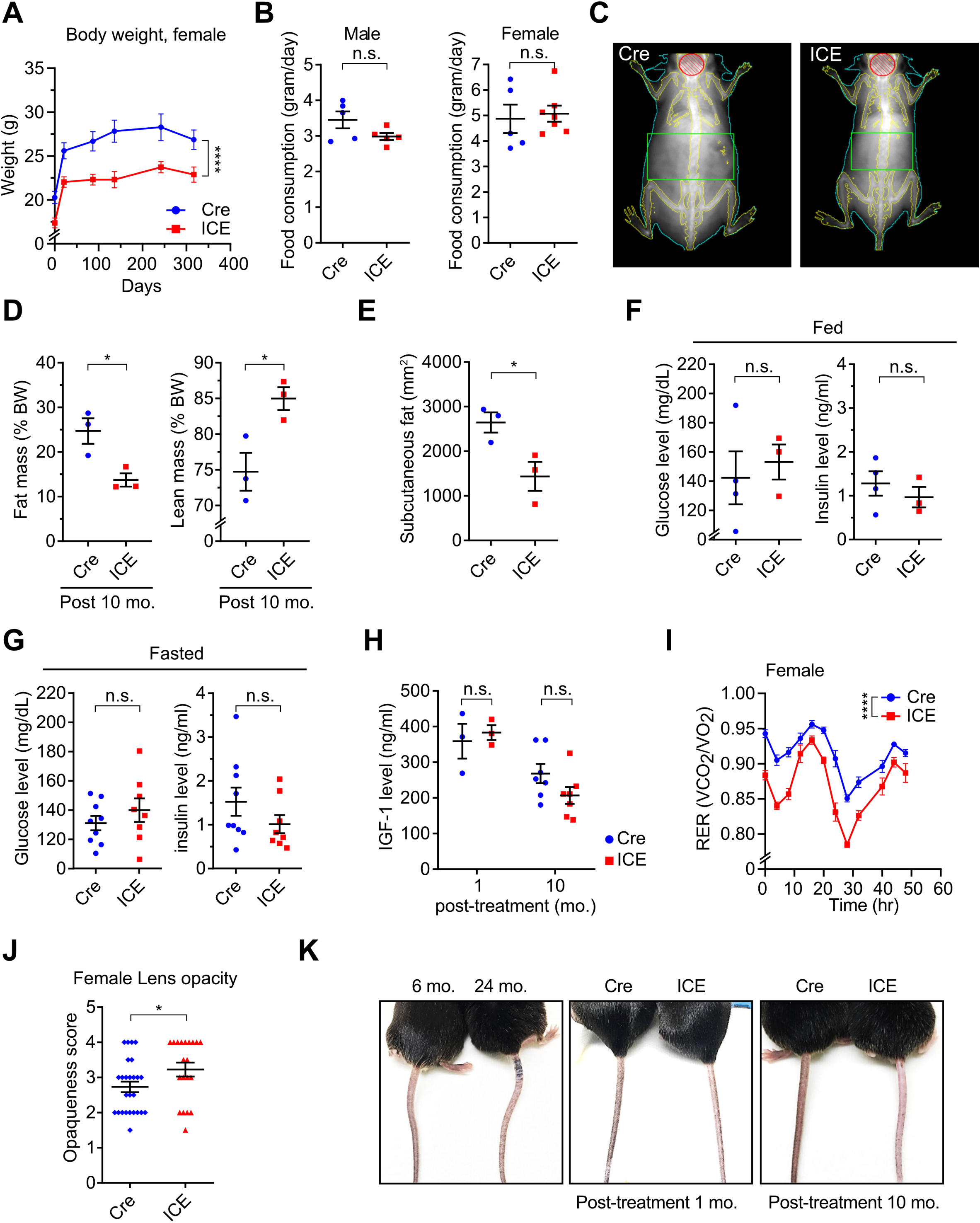
ICE Mice Phenocopy Normal Aging, Related to Figure 3. (A) Body weights of female Cre and ICE mice 10-month post-treatment. Repeated measures one-way ANOVA. (B) Food consumption of post-treated Cre and ICE mice. Two-tailed Student’s *t* test. (C) Representative images of Cre and ICE mice from Dual Energy X-ray Absorptiometry (DEXA) 10-month post-treatment. (D) Body mass of 10-month post-treated Cre and ICE mice. Two-tailed Student’s *t* test. (E) Subcutaneous fat thickness of 10-month post-treated Cre and ICE mice. Two-tailed Student’s *t* test. (F and G) Blood glucose levels of 10-month post-treated Cre and ICE mice in the fed or fasted state. Two-tailed Student’s *t* test. (H) IGF-1 levels in 10-month post-treated Cre and ICE mice. Two-way ANOVA-Bonferroni. (I) Respiratory Exchange Rate (RER) of female Cre and ICE mice. Repeated measures one-way ANOVA. (J) Lens opacity of 10-month post-treated female Cre and ICE mice. Two-tailed Student’s *t* test. (K) Representative tail pigmentation of wild type, Cre and ICE and mice. Data are mean ± SEM. n.s.: p > 0.05; *p < 0.05, ****p< 0.0001.

**Figure S5.**
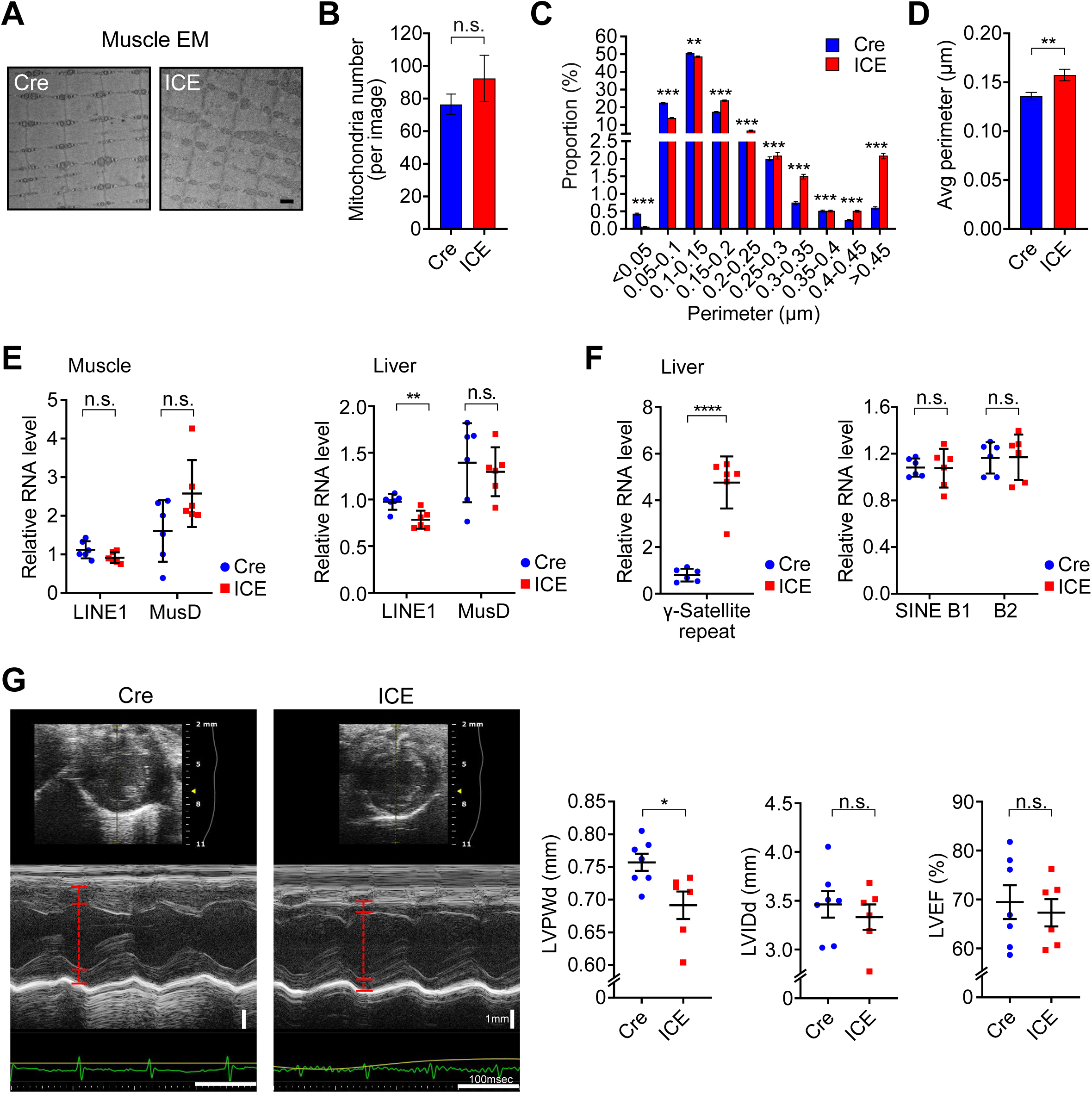
Progeroid Features of ICE Mouse Muscle, Related to Figure 5. (A-D) Mitochondria number (A and B) and mitochondrial perimeter (A, B and D) of 10-month post-treated Cre and ICE muscle determined with electron microscopy. Two-tailed Student’s *t* test. (E and F) Quantification of RNA from repetitive DNA elements in gastrocnemius muscle and liver from 10-month post-treated Cre and ICE mice. Two-tailed Student’s *t* test. (G) Echocardiogram of 10-month post-treated Cre and ICE mice. LVPWd, Left ventricular posterior wall thickness at end-diastole mm. LVIDd, Left ventricular internal diameter in diastole; LVPWd, Left ventricular posterior wall in diastole; LVEF, Left ventricular ejection fraction. Two-tailed Student’s *t* test. Data are mean ± SEM. n.s.: p > 0.05; *p < 0.05; **p < 0.01; ***p< 0.001, ****p< 0.0001.

**Figure S6.**
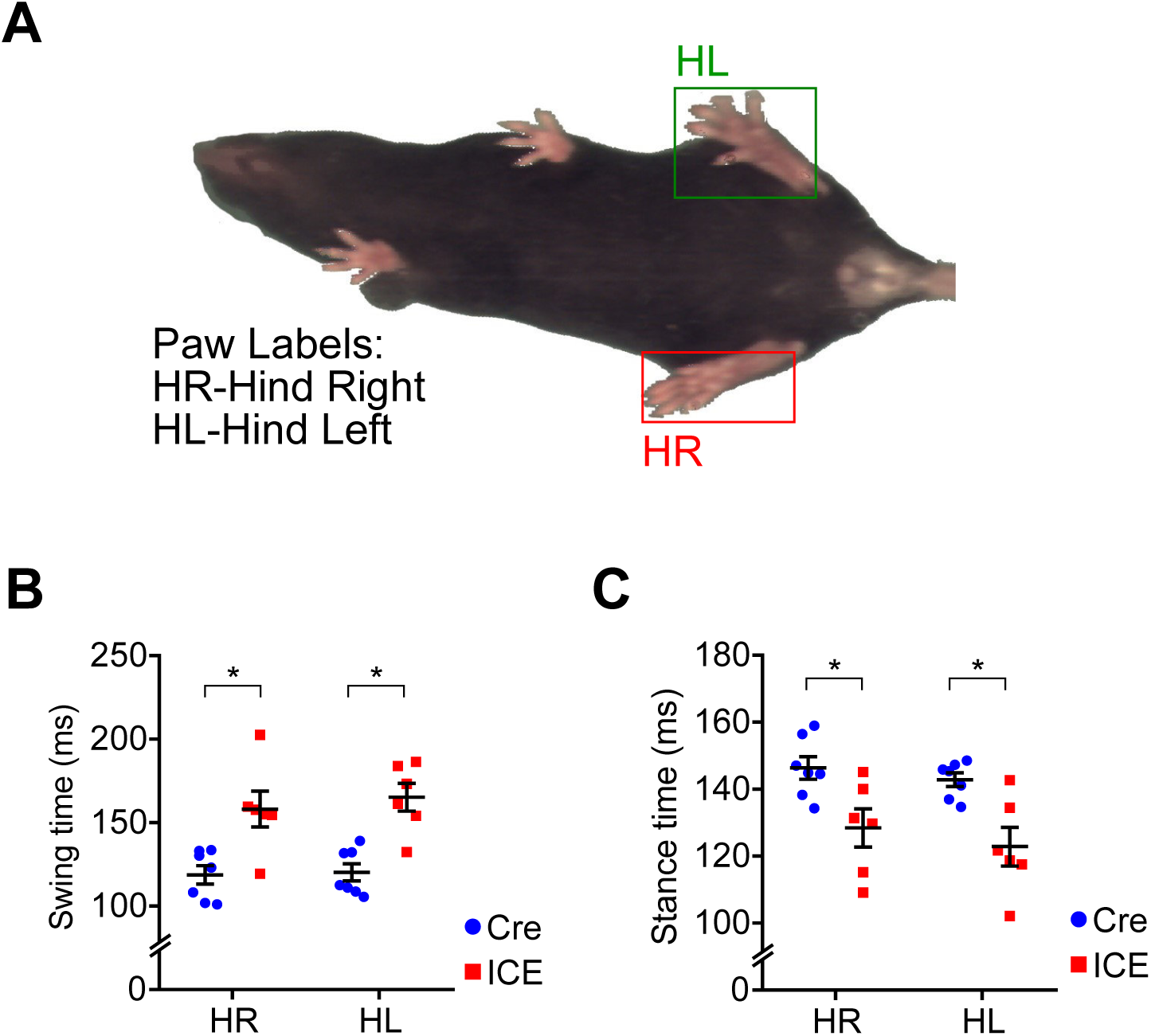
Impaired Gait Coordination in ICE Mice, Related to Figure 6. (A-C) Gait analysis of Cre and ICE mice. Two-tailed Student’s *t* test. Data are mean ± SEM. *p < 0.05.

**Figure S7.**
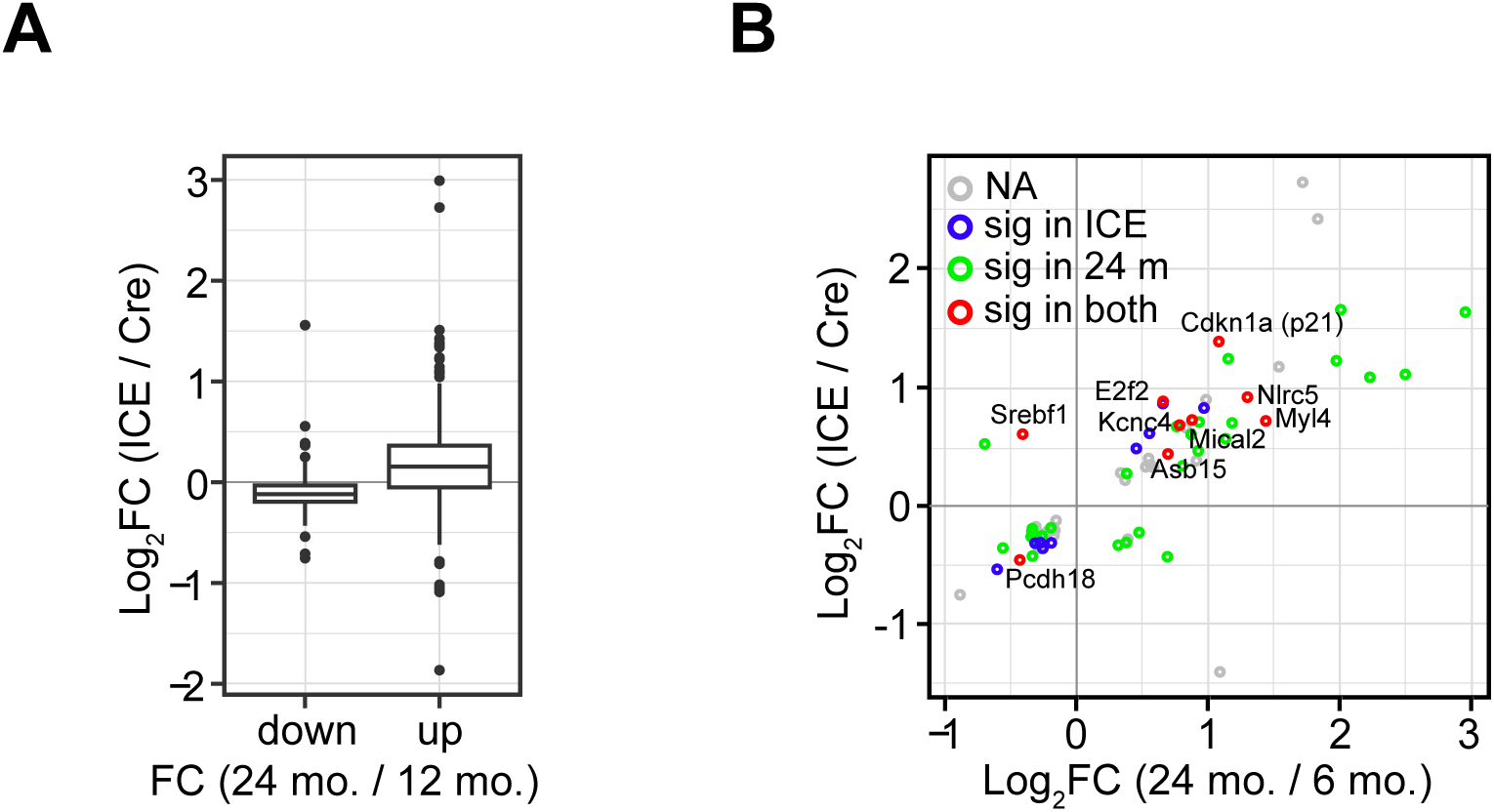
ICE Mice Mimic Transcriptomic Changes of Wild Type Old Mice, Related to Figure 7. (A) Scatter plot of genes significantly altered (p<0.01) in 10-month post-treated ICE mice or wild type 24 month-old mice. (B) Fold change of altered genes (padj<0.05, 12mo. vs 24mo.) in 10-month post-treated ICE mice.

